# Humoral correlate of vaccine-mediated protection from tuberculosis identified in humans and nonhuman primates

**DOI:** 10.1101/2024.12.05.627012

**Authors:** Natasha S. Kelkar, Nicholas C. Curtis, Urjeet S. Khanwalkar, Olivia G. Palmer, Tim P. Lahey, Wendy Wieland-Alter, Jason E. Stout, Erica C. Larson, Solomon Jauro, Charles A. Scanga, Patricia A. Darrah, Mario Roederer, Robert A. Seder, C. Fordham von Reyn, Jiwon Lee, Margaret E. Ackerman

## Abstract

Development of an effective tuberculosis (TB) vaccine has been challenged by incomplete understanding of specific factors that provide protection against *Mycobacterium tuberculosis* (Mtb) and the lack of a known correlate of protection (CoP). Using a combination of samples from a successful vaccine efficacy trial and studies of Bacille Calmette-Guerin (BCG)-immunized humans and nonhuman primates (NHP), we identify a humoral CoP that translates across species and vaccine regimens. In a human efficacy study, antibodies specific to the DarDar vaccine strain (*M. obuense*) sonicate (MOS) correlate with protection from microbiologically confirmed TB. In humans, antibodies to MOS also scale with vaccine dose, are elicited by BCG vaccination, and demonstrate cross-reactivity with Mtb. In NHP, MOS-specific antibodies scale with dose and correlate with TB protection mediated by BCG vaccination. Collectively, this study reports a novel humoral CoP and specific antigenic targets that may be relevant to achieving vaccine-mediated protection from TB.

**Summary:** An effective vaccine is crucial to achieving control of tuberculosis (TB), but the lack of a reliable correlate of protection (CoP) presents a major hurdle to TB vaccine development. Kelkar et al. report a novel antibody CoP that generalizes across vaccines and identifies antigenic targets that may be relevant to achieving vaccine-mediated protection from TB.

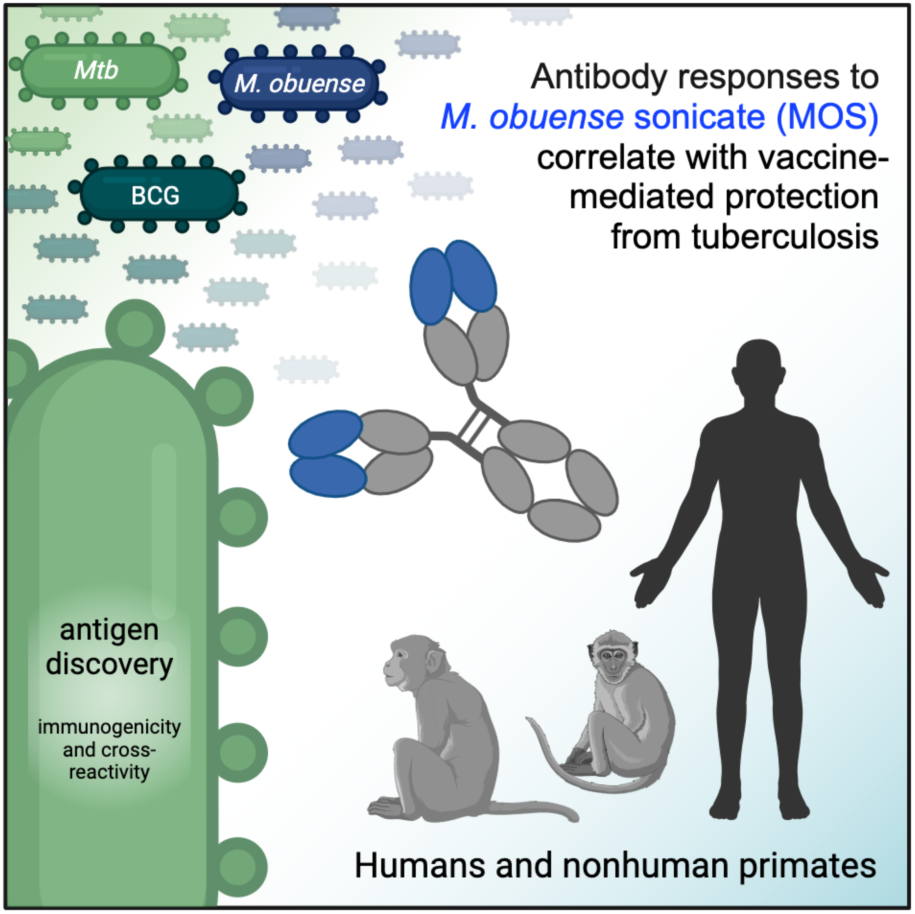

## Introduction

Tuberculosis (TB) is the leading infectious disease cause of death globally. Based on the current annual TB death rate of 2%, predicted TB mortality from 2020 to 2050 is estimated at 31.8 million deaths, corresponding to an economic loss of 17.5 trillion USD^1^. The Bacillus Calmette-Guérin (BCG) vaccine, introduced in 1921, is the only effective vaccine currently available. BCG, which also reduces all-cause mortality^2^, is an attenuated strain of *Mycobacterium bovis*, the etiological agent of TB in cattle. However, a birth dose of this vaccine has modest efficacy that wanes after 10-15 years^3,4^, motivating the development of booster strategies for adolescents and adults immunized at birth^5^.

Unfortunately, TB vaccine development has been slowed by the absence of known human correlates of protection (CoP) after vaccination with BCG or other candidate vaccines^3,6^. Identification and validation of CoP could meaningfully decrease the time and cost of early development assessments of candidate vaccine regimens^7^ and contribute to successful development and deployment of improved TB vaccines, which are crucial to global TB control^8^. With this goal in mind, we profiled humoral immune responses to inactivated whole cell *Mycobacterium obuense* (*M. obuense*) SRL172 vaccine, a post-BCG booster vaccine candidate which conferred protection from culture-confirmed TB in a randomized, double-blind, placebo-controlled phase III clinical trial (DarDar) in Tanzania [NCT00052195]^9,10^. Building on phase I and phase II studies in Finland and Zambia confirming safety and immunogenicity in people living with HIV (PLWH) with prior BCG vaccination ^11,12^, the phase III DarDar trial was stopped early for statistically significant vaccine efficacy of 39% for the secondary endpoint of preventing culture-confirmed TB, and thus represents an important step toward improved prevention of TB^9^. SRL172 vaccine elicited cellular and humoral immune responses, but limited testing suggested these were of low magnitude and no CoP was identified^13^, leaving markers and mechanisms of protection undefined. Archived samples from this trial provide an unprecedented opportunity to interrogate immune correlates of vaccine-mediated protection from TB that complement immunogenicity studies of other contemporary TB vaccine candidates.

While there is much evidence to support the mechanistic relevance of cellular immune responses to protection from TB in humans and preclinical models, these responses are considerably more complex to measure than the humoral responses prevalent among CoP known for other protective vaccines. Furthermore, there is ample evidence to support seeking humoral correlates of vaccine-mediated protection from TB. Humans diagnosed with active or latent TB exhibit differential serum antibody composition and function^14–17^, and serum IgG from *Mycobacterium tuberculosis* (Mtb)-exposed healthcare workers contains Mtb surface-specific antibodies that inhibit Mtb growth *in vitro* and reduce bacterial burden in mouse challenge studies^18^. Antibody depletion and passive immunization studies of monoclonal antibodies in mice further confirm the potential for mechanistic humoral contributions to protection from TB^19–32^, and BCG-vaccinated macaques raise humoral responses that are associated with prevention of TB infection^33–35^.

Humoral responses may be especially relevant in the context of diminished T cell counts and function, such as from normal human aging or co-infection with Mtb and HIV, a major global syndemic. The two pathogens potentiate one another^36^, with HIV infection rates rising as protection from TB afforded by BCG immunization at birth wanes. Indeed, TB disease is the most common opportunistic infection causing mortality in PLWH^37^, accounting for roughly 30% of world-wide AIDS-related deaths^38,39^. Thus, the development of a vaccine, like SRL172, that is safe and effective for PLWH is a leading global health priority^40,41^. To this end, while not directly translatable, pre-clinical models of i.v administration of BCG^35,42–44^, which has demonstrated near complete protection in nonhuman primate (NHP) models, highlight a new route by which a century-old vaccine may yet influence global health. Here, we leverage archived samples from these and other studies to survey for humoral CoP to aid and guide future TB vaccine development.

## Results

### Vaccination with SRL172 induces *M. obuense* sonicate (MOS)-specific antibodies

Participants in the DarDar trial included PLWH with a CD4 count of at least 200 cells/μL and a BCG scar who received either a five-dose series of 1 mg inactivated *M. obuense* SRL172 or borate-buffered isotonic saline intradermally (**Figure 1A**)^9^. Serum samples from trial participants, who were screened for TB routinely every three months, were selected using a case-control design and evaluated using systems serology approaches. Participants who developed definite TB (cases) following completion of the five-dose series in both vaccine and placebo arms were matched based on age, tuberculin skin test results, and prior TB status with control subjects who did not develop definite TB after vaccination (**Supplemental Table 1**).

**Fig. 1:**
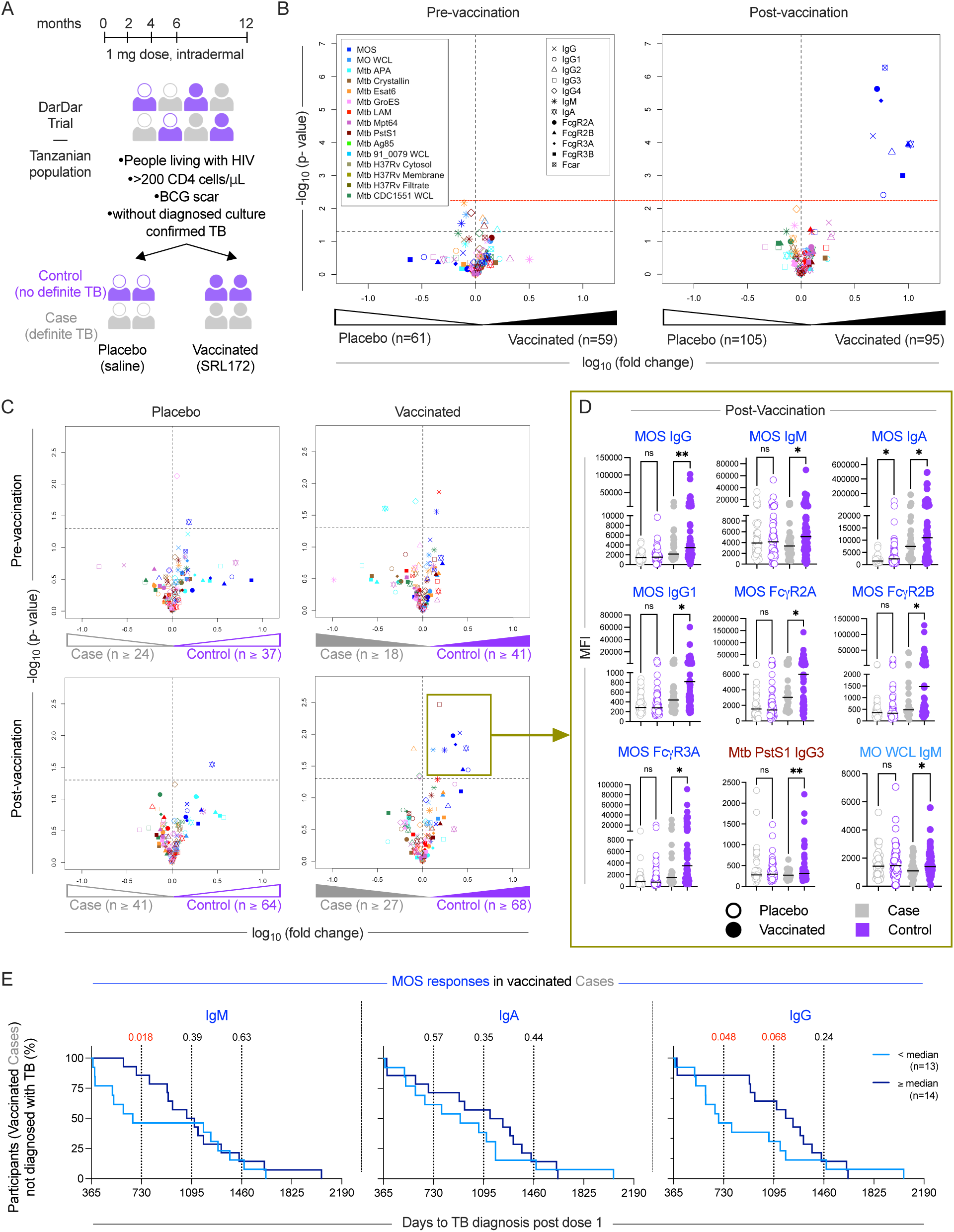
MOS-specific antibodies correlate with protection from disease in the DarDar trial. **A.** Schematic of the DarDar vaccine trial conducted in Dar es Salaam, Tanzania. Participants were randomized to receive five intradermal immunizations over one year with 1 mg dose of the vaccine (SRL172 vaccine) or placebo (borate buffered isotonic saline). **B-C.** Volcano plots depicting the magnitude (fold change) and statistical significance (Welch’s t-test) of differences in each measured humoral immune response feature detected in serum samples from placebo and vaccinated subjects at baseline (pre-vaccination, left) and following immunization (post-vaccination, right) (**B**), and between Cases and Controls in measured immune features at pre-vaccination (top) and post-vaccination (bottom) timepoints and in placebo (left) and vaccinated (right) participants (**C**). Symbol color indicates antigen specificity, while shape indicates the antibody Fc domain characteristic for each response feature measured. Dotted horizontal lines indicate unadjusted p = 0.05 as determined by Welch’s t-test in black. For panel B, the false positive significance threshold set by differences observed at baseline is indicated in red. For panel C, adjusted significance values are presented in **Supplemental Table 4**. **D.** Box plots comparing the levels of select (gold box) immune features at post-vaccination timepoint amongst cases (gray) and controls (purple) in placebo (hollow circles) and vaccine (solid circles) recipients. Statistical significance was determined by Unpaired t-test with Welch’s correction (**p<0.01, *p<0.05, ns p ≥ 0.05). Red bar indicates median. **E.** Kaplan-Meier curves depicting diagnosis of TB over time in vaccinated cases for high (≥ median, dark blue) versus low (< median, light blue) MOS-specific IgM (left), IgA (center) and IgG (right) responders. Statistical significance was evaluated at years 1, 2, and 3 (vertical lines) post final vaccine dose using log-rank (Mantel-Cox) test; values <0.1 are indicated in red. Subject number (n) is indicated in each figure panel, with the exception of panel D, for which placebo case (n=41), placebo control (n=64), vaccinated case (n=27), vaccinated control (n=68).

We evaluated antibody responses in blinded serum samples drawn from participants at the pre- (n=120) and post- (n=200) immunization timepoints. Antigen-specific IgA, IgM, IgG and IgG subtypes (IgG1, IgG2, IgG3, IgG4) responses were evaluated^45,46^. Characterization extended beyond isotypes and subclasses to include propensity to bind Fc receptors (FcγR2A, FcγR2B, FcγR3A, FcγR3B and FcαR). The panel of antigens consisted of sonicate derived from the SRL172 vaccine strain *M. obuense* grown on agar (MOS); whole cell lysate of vaccine strain *M. obuense* grown in broth (MO WCL); Mtb virulence factor early secreted antigen target 6kDa (ESAT6), alanine- and proline-rich antigenic protein (APA), lipoarabinomannan (LAM), α-crystallin, PstS1, the major culture filtrate protein Mpt64, the heat shock protein GroES, evasion factor antigen 85 complex; WCL from Mtb strain 91_0079; Mtb strain CDC1551 WCL, cytosolic, and cell membrane fractions, as well as culture filtrates of Mtb strain H37Rv (**Supplemental Table 2**).

Whereas a few nominal differences of small magnitude and low confidence in measured antibody responses between placebo- and SRL172 recipients were observed at baseline (**Figure 1B)**, none were significant following multiple hypothesis correction (**Supplemental Figure 1, Supplemental Table 3**). In contrast, robust and confident false discovery rate (FDR)-adjusted differences in MOS-specific IgG, IgG1, IgG2, IgA and binding to FcαR, FcγR2A, FcγR2B, FcγR3A, and FcγR3B were observed in SRL172 as compared to placebo recipients post-immunization (**Figure 1B**, **Supplemental Figures 1-2, Supplemental Table 3**). Overall, this analysis demonstrated induction of diverse isotypes and subclasses of antibodies specific for the vaccine strain.

### MOS-specific antibodies correlate with protection in the DarDar trial

To identify a CoP, we compared humoral responses over time in placebo- and vaccine-recipients in participants who did or did not develop TB disease using a case-control design. While TB disease status groups did not differ in CD4 T cell counts or HIV-1 viral load at baseline (**Supplemental Figure 3**), we observed elevated levels of MOS-specific IgM, IgA, IgG, IgG1, and FcγR2A-, FcγR2B-, FcγR3A-binding antibody responses in vaccinated controls compared to vaccinated TB disease cases in cross-sectional analysis between groups (**Figure 1C**, **1D**). Longitudinal analyses demonstrated robust induction of these humoral responses among vaccine recipients that did not experience TB disease (**Supplemental Figure 4**). While Mtb PstS1-specific IgG3 responses were also nominally elevated in controls (**Figure 1C**), differences in magnitude were small (**Figure 1D**). IgM responses to MO WCL were also elevated in vaccinated controls as compared to cases. Among the responses induced by SRL172 (**Supplemental Figure 1, Supplemental Table 1**), levels of MOS-specific IgG, IgG1, IgA, FcγR2A, FcγR2B, FcγR3A binding antibodies were elevated in vaccinated controls relative tovaccinated cases following multiple hypothesis correction (FDR-adjusted p <0.01) (**Supplemental Table 4**), demonstrating that MOS-specific antibodies are a CoP in the DarDar trial. In contrast, relatively few nominal differences between cases and controls were noted in pre-immunization response profiles or among placebo-recipients at either timepoint. Intriguingly, however, MOS-specific IgA was elevated at both pre- and post-immunization timepoints in placebo controls as compared to cases, suggesting that it may be a marker of pre-existing immunity to TB (**Figure 1C**, **1D, Supplemental Figure 4**).

### Differential risk is associated with MOS-specific antibody response magnitude among breakthrough TB disease cases in vaccine recipients

To further evaluate the relevance of MOS-specific antibodies as CoPs, we next tested relationships between time to TB diagnosis and MOS-specific antibody response magnitude within the subset of participants who ultimately developed definite TB. TB cases in either the placebo or SRL172 recipient groups were each classified as ‘low’ or ‘high’ responders based on the magnitude of MOS-specific IgM, IgA, or IgG responses post-vaccination. Among vaccine but not placebo recipients who were diagnosed with definite TB, those with IgM and IgG responses to MOS at or above the group median showed significantly decreased risk of disease early in the follow up period (**Figure 1E, Supplemental Figure 5**). These results demonstrate that MOS IgG and IgM responses are also correlates of reduced risk of early disease diagnosis in breakthrough cases, and a quantitative relationship exists between response magnitude and time to disease.

### Immunogenicity of *M. obuense* across TB vaccines

Preparations derived from both agar- (SRL172) and broth-grown (DAR-901^47^) culture of *M. obuense*, have been in development as a post-BCG booster vaccine. DAR-901 vaccine, which has completed phase I and II safety and immunogenicity trials in participants with and without HIV infection^47,48^, was produced using a broth-based manufacturing process from the *M. obuense* SRL172 master cell bank to improve scalability (**Figure 2A**). To assess immunogenicity of the DAR-901 vaccine candidate, shown in a murine model to provide protection against TB challenge superior to BCG^49^, serum drawn from 58 healthy adult participants with prior BCG vaccination and negative TB IFN-γ release assay who received either three intradermal administrations of DAR-901 at 0.1 mg (n=10), 0.3 mg (n=10) or 1.0 mg (n=20) dose, or three doses of saline placebo (n=9), or two doses of saline and one dose of BCG (n=9) at 0, 2, and 4 months (**Supplemental Table 5**) were evaluated. Sera from US-based participants in the two higher dose groups of the phase I dose escalation trial (NCT02712424) of DAR-901^47^ demonstrated recognition of MOS (**Figure 2B**) at similar or higher levels than in those who received SRL172 in the DarDar trial (**Supplemental Figure 6**). Longitudinal profiling of samples prior to immunization (Pre) or after dose 3 (Post) demonstrated dose-dependent increases in MOS-specific IgG (**Figure 2B**). Although smaller in magnitude, MOS-specific IgG was also seen following a single immunization with BCG, demonstrating that antibodies induced or boosted by BCG cross-react with antigens in MOS. In contrast, there was no increase in MOS-specific IgG in the placebo group. Neither DAR-901 nor placebo induced antibodies that cross-reacted with Mtb CDC1551 WCL, whereas immunization with BCG did (**Figure 2B**). These results demonstrate that additional TB vaccines, including the efficacious BCG vaccine, induce MOS-specific antibodies in humans. Further, mycobacteria can induce responses with varying cross-reactivity profiles. The variable magnitude of increases in Mtb CDC1551-specific antibodies following BCG immunization than DAR-901 may reflect differing degrees of antigenic and immunogenic similarity among strains. Evidence of cross-reactivity between strains notwithstanding, the detection of induced cross-reactivity from re-immunization of adults does not inform on MOS-specific antibodies as a CoP for childhood doses of BCG. Additionally, because the tested samples were from a phase 1 dose-escalation study, infection status outcomes were not available, and relationships between MOS-specific antibodies and TB infection or disease could not be evaluated.

**Fig. 2:**
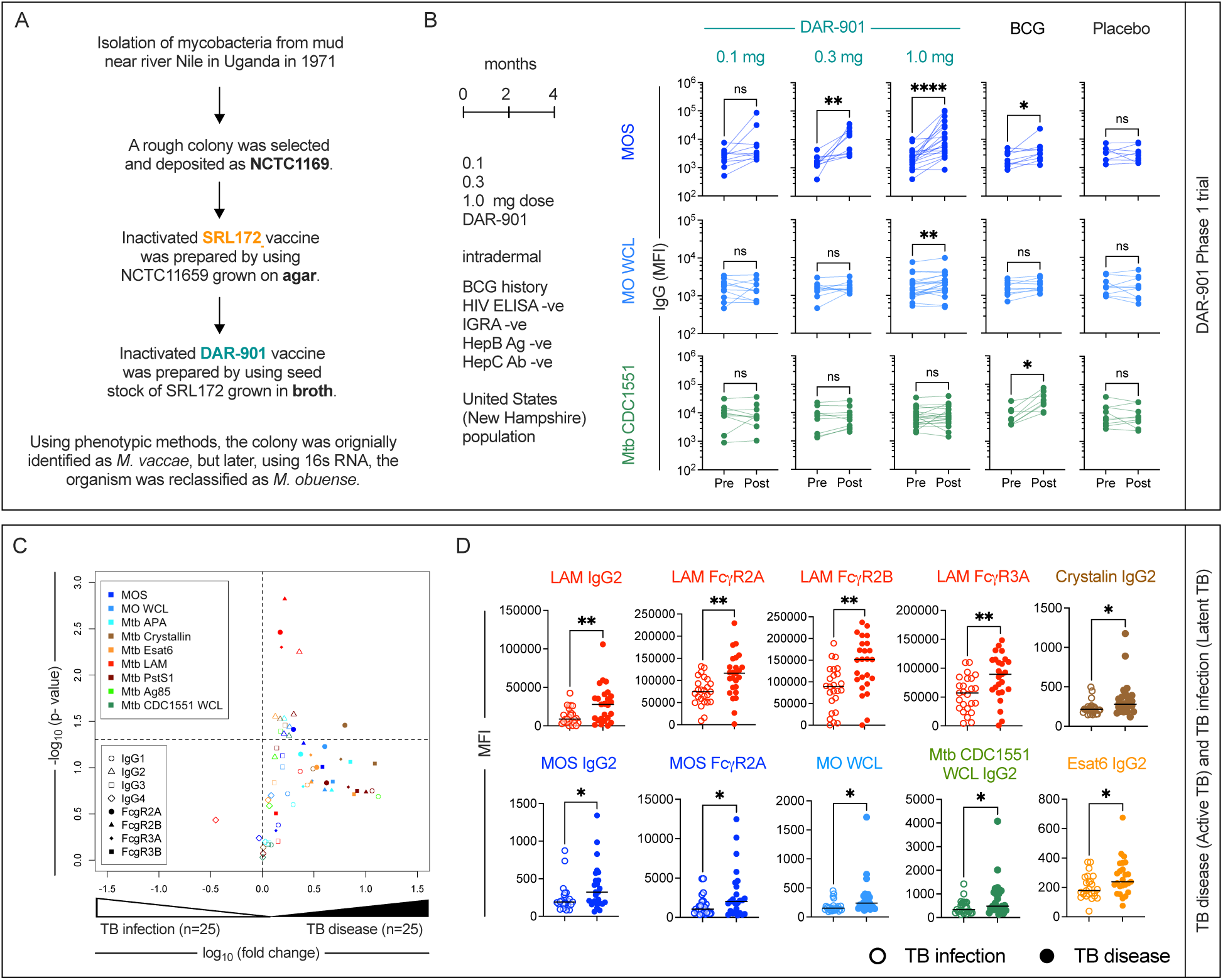
Cross-reactivity antibodies induced by DAR-901 vaccine, BCG vaccination, and Mtb infection to MOS. **A.** History of preparation of DAR-901 vaccine. **B.** Longitudinal profiling of IgG specific for MOS (top), MO WCL (middle), and Mtb CDC1551 WCL (bottom) profiling for US participants of DAR-901 trial Phase 1 trial who were administered either three doses of DAR-901 vaccine at 0.1 (n=10), 0.3 (n=10), or 1.0 mg (n=20) doses, three doses of saline (Placebo, n=9), or two doses of saline and one dose of BCG vaccine (BCG, n=9) at 0, 2, and 4 months intradermally. Statistical analysis was performed by Wilcoxon matched paired sign rank test (****p<0.0001, ***p<0.001, **p<0.01, *p<0.05). **C.** Volcano plots depicting the magnitude (fold change) and statistical significance (Welch’s t-test) of differences in measured immune features in individuals diagnosed with tuberculosis infection (n=25) or disease (n=25). Dotted horizontal line indicated unadjusted p value of 0.05. Color indicates antigen specificity, while shape indicates the antibody Fc domain characteristic for each response feature measured. **D.** Box plots comparing the levels of select immune features in subjects with Latent (hollow circle) or Active (solid circle) tuberculosis. Statistical significance was determined by Welch’s t-test (**p<0.01, *p<0.05). Bar indicates median.

### MOS-cross-reactive IgG2 is associated with active TB disease

While evaluation following BCG immunization suggested that cross-strain responses could be induced, there was little evidence that *M. obuense* SRL172 induced antibodies that recognized antigens derived from the limited number of Mtb strains in multiplex assay testing (**Figure 1B**). To address whether antibodies associated with Mtb infection could cross-react with MOS, we evaluated responses in a cohort of individuals diagnosed with either active TB disease (n=25) or latent Mtb infection (n=25) (**Supplemental Table 6**).

Consistent with prior reports^17,50^, we observed generally elevated humoral responses in individuals with active TB disease (**Figure 2C**). Specifically, we observed higher LAM-specific responses, specifically IgG2 and binding to FcγR2A, FcγR2B and FcγR3A in individuals with active TB disease (**Figure 2D**). Multiple Mtb antigens were targeted by elevated levels of IgG2 in individuals with active TB disease, including ESAT6, α-crystallin, APA, PstS1, and Mtb CDC1551 WCL. Lastly, MOS- and MO WCL-specific IgG2 responses were also elevated, the former also showing elevated binding to FcγR2A, which was evaluated with the H131 allotype capable of binding human IgG2 (**Figure 2D**). Skewing toward IgG2 across a range of antigen-specificities suggests differential regulation of plasmablasts secreting this subclass in the context of active disease, and elevated MOS-specific IgG2 suggests boosting of cross-reactive antibodies during active TB infection and disease despite strain differences. While supporting shared immunogenicity and antigenicity between disease-causing strains and *M. obuense*, these data do not provide evidence for or against correlative relationships between MOS-specific antibodies and vaccine-mediated protection.

### BCG vaccination elicits MOS-specific antibodies in SIV-infected and SIV-naïve cynomolgus macaques

To explore whether cross-reactivity between MOS- and protective TB-specific humoral responses exists in other contexts, we measured relative titers of MOS- and Mtb CDC1551 WCL-specific IgG antibodies in plasma samples acquired from two NHP species, with and without SIV infection, and immunized with BCG at various doses and routes (**Supplemental Table 7**).

Paralleling the DarDar trial assessment of MOS- and TB-specific humoral responses in PLWH, *Macaca fascicularis* (cynomolgus macaques) infected with simian immunodeficiency virus (SIV) strain SIVmac239 were immunized with BCG either intradermally (i.d.) with 5×10^5^ colony forming units (CFU) (n=6) or i.v. with 5×10^7^ CFU followed three weeks later by treatment with isoniazid, rifampicin, ethambutol (HRE) to prevent disease caused by BCG (n=5)^43^ (**Figure 3A**). Plasma samples were collected pre- and one month post-vaccination. Whereas four of five animals receiving i.v. BCG, which confers greater protection than i.d. BCG in SIV-naïve animals^35^, showed increased MOS- and Mtb CDC1551 WCL-specific IgG, such responses were not elicited following i.d. BCG vaccination (**Figure 3A**).

**Fig. 3:**
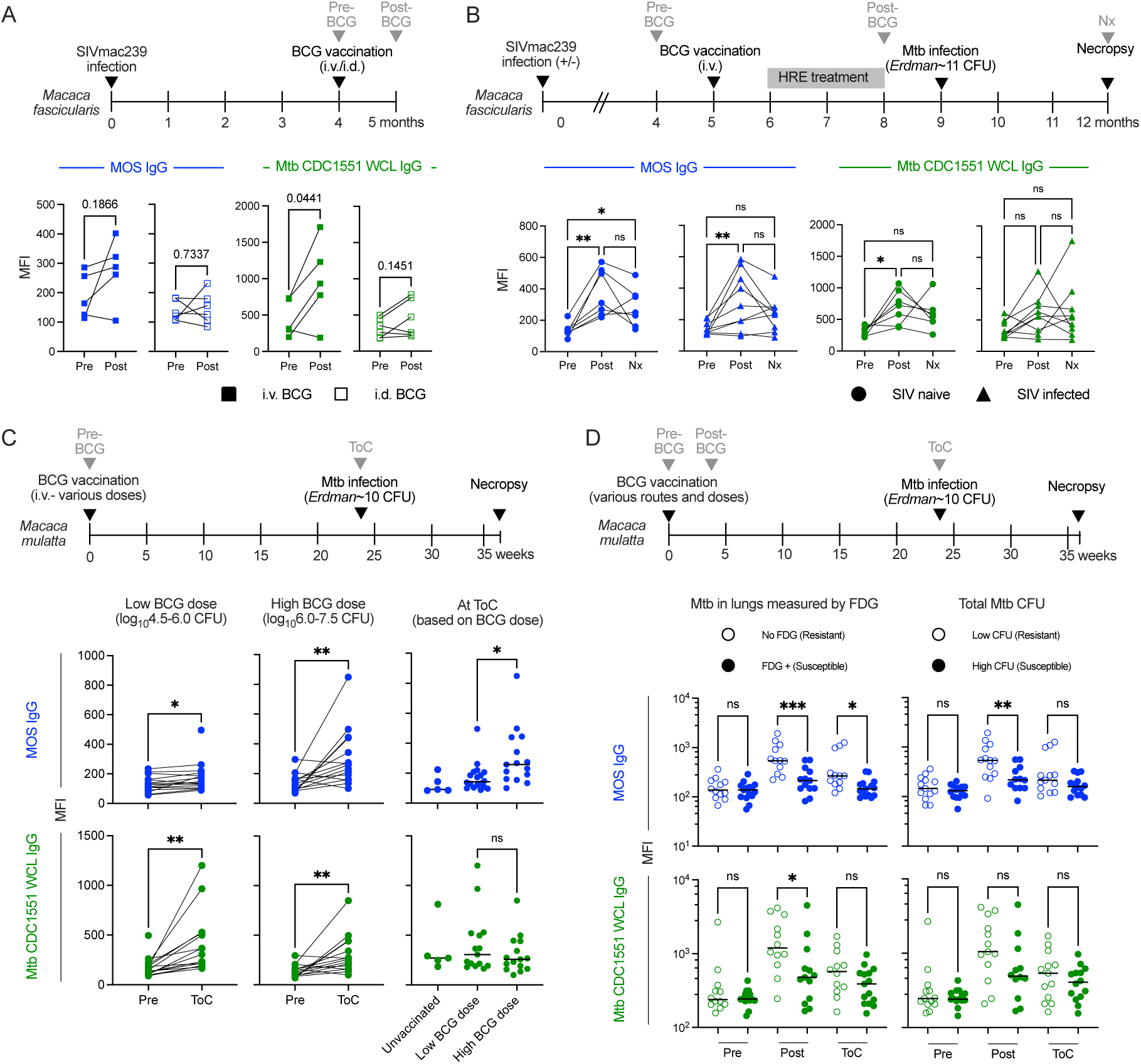
MOS-specific IgG responses are induced by BCG immunization in NHP and are associated with vaccine-mediated protection. **A.** Longitudinal profiling at pre- and post-vaccination timepoints of MOS- (blue) and Mtb (CDC1551) WCL- (green) specific IgG responses in SIV-infected *M. fascicularis* administered i.d. (n=6) or i.v. BCG (n=5) and HRE (Isoniazid, Rifampin, Ethambutol) therapy. Statistical significance was determined by paired t-test. **B.** Longitudinal profiling at pre-vaccination, post-vaccination and necropsy (Nx) timepoints of MOS-and Mtb WCL-specific IgG responses in SIV-infected (n=9) and naïve (n=7) *M. fascicularis* who were administered i.v. BCG and protected from Mtb challenge. Statistical significance was determined by paired one-way ANOVA. **C.** Longitudinal profiling at pre-vaccination and time of Mtb challenge (ToC) in *M. mulatta* vaccinated with low (left, n=18) and high dose i.v. BCG (center, n=16). Statistical significance was determined by paired t-test. MOS- and Mtb WCL-specific IgG responses at ToC by BCG vaccine history (right). Statistical significance was determined by Welch’s t test. **D.** Longitudinal profiling of MOS- and Mtb WCL-specific IgG responses in *M. mulatta* (n=28) by Mtb infection burden determined by 2-deoxy-2-(^18^F) fluorodeoxyglucose (FDG) imaging (left) and by colony forming units (CFU) of Mtb in lungs (right). Statistical significance was determined by one-way ANOVA. Significance is indicated as: ***p<0.001, **p<0.01, *p<0.05, ns p≥0.05.

The capacity of i.v. BCG to mediate protection against TB in SIV-infected animals was directly addressed in another study in which cynomolgus macaques with or without prior SIVmac239 infection were immunized with i.v. BCG (**Figure 3B**). To preclude symptomatic disseminated BCG, animals received a two-month regimen of HRE three weeks after vaccination that was discontinued one month prior to challenge with low dose Mtb Erdman (∼11 CFU) via bronchoscopic instillation^42^ (**Figure 3B**). MOS- and Mtb CDC1551 WCL-specific IgG responses were evaluated in the seven of seven SIV-naïve and nine of twelve SIV-infected animals that were protected from Mtb challenge by i.v. BCG^42^. Whereas BCG vaccination induced MOS-specific IgG irrespective of SIV infection status, Mtb CDC1551 WCL-specific IgG was induced only in SIV naïve animals (**Figure 3B**), suggesting that at least for these isolate preparations, SIV infection impaired induction of cross-reactive Mtb- but not MOS-specific humoral responses.

### MOS-specific IgG is a CoP in BCG-vaccinated rhesus macaques

To validate its observation as a CoP in humans and to assess the generalizability of the association between anti-MOS antibody responses and protection from TB disease, we next evaluated the effect of i.v. BCG on protection from TB in *Macaca mulatta* (rhesus macaques)^44^.

Rhesus macaques were immunized with i.v. BCG (4.5-7.5 log_10_ CFU) and challenged with low-dose Mtb Erdman at ∼six months post-vaccination (**Figure 3C**). Both MOS- and Mtb CDC1551 WCL-specific IgG (**Figure 3C**) but not IgM or IgA (**Supplemental Figure 7A-C**) responses were observed following vaccination with either low or high dose BCG. At the time of Mtb challenge, higher MOS- but not Mtb CDC 1551 WCL-specific IgG was detected in animals that received high (6.0-7.5 log_10_CFU) as compared to low (4.5-6.0 log_10_CFU) dose BCG, demonstrating a dose-response relationship between antibody generation and BCG vaccine dose, which is known to influence protection from TB^44^.

Next, we analyzed plasma from a study of the effect of route and dose of BCG vaccination on protection^35^. Rhesus macaques (n=25) were vaccinated i.d. with high or low dose BCG, or i.v. with high dose BCG prior to bronchoscopic challenge with Mtb Erdman (∼10 CFU). Challenge outcomes were measured by evaluating inflammation in the lungs with PET-CT imaging using 2-deoxy-2-(^18^F) fluorodeoxyglucose (FDG), as well as by determining total Mtb CFU in lung at necropsy. Animals were split by median into two groups: Mtb resistant (low FDG activity or low Mtb CFU at necropsy) or susceptible (>300 FDG activity or high Mtb CFU at necropsy). Resistant animals, as defined by FDG activity, comprised mostly of i.v. BCG recipients, demonstrated higher MOS- and Mtb CDC 1551 WCL-specific IgG at the post-vaccination as compared to the susceptible group, which was comprised mostly of i.d. low dose BCG recipients (**Figure 3D**), linking the magnitude of MOS-specific IgG to disease resistance. This difference was maintained at the time of challenge for MOS- but not Mtb CDC 1551 WCL-specific IgG. Supporting the relevance of this relationship, no differences in antibody levels were observed at the pre-vaccination timepoint. Considering the CFU endpoint, only MOS-specific IgG at the post-immunization timepoint served as a correlate of disease (**Figure 3D**). TB resistant macaques also demonstrated higher MOS-specific IgA responses post-vaccination as compared to the susceptible animals (**Supplemental Figure 7D**). Further, despite the observation that MOS-specific IgG responses post-vaccination did not always differ in magnitude in association with route or dose (**Supplementary Figure 8**), the magnitude of antibody responses to MOS antigens did differ in association with the degree of disease resistance across all animals for both FDG (R_s_=-0.57; p=0.0029) and CFU (R_s_=-0.55; p=0.0047), as well as within the i.d. high dose group for CFU (R_s_=-0.83; p=0.015) at the post-immunization timepoint (**Supplementary Figure 9**). Responses to Mtb CDC 1551 WCL did not show such associations.

Overall, results from these diverse NHP studies demonstrate that, as in humans, BCG vaccination induces antibodies that cross-react with MOS. Elicitation of MOS-specific IgG responses was BCG dose-dependent, observed in two different NHP species, including animals infected with SIV, and, though confounded by differences in vaccine route and dose, correlated with infection outcomes. This independent validation bolsters the potentially broader relevance of MOS-specific antibodies as a CoP against Mtb.

### Identification of immunogenic MOS antigens

To identify specific components of MOS recognized by antibodies raised in vaccinated participants in the DarDar trial, we used affinity purification to isolate MOS-specific antibodies post-immunization from a selected subset high IgG responders including ten vaccinated participants who did not develop TB disease (controls) and three vaccinated participants who did develop TB disease (cases) (**Supplemental Figure 10A**). Pooled total serum IgG from the pre-vaccination timepoint for seven of the ten vaccinated controls from whom MOS-specific Ig was isolated was used as a control. These antibody preparations were then used to affinity purify immunogenic components of MOS, which were then identified using mass spectrometry (**Supplemental Figure 10B**).

A total of 1,087 proteins were identified in MOS (**Figure 4A**). Nine of these proteins (from ketol-acid reductoisomerase (NADP(+)), PspA/IM30 Chaperonin GroEL, ABC transporter substrate binding protein, cytosol aminopeptidase, glyceraldehyde-3-phosphate dehydrogenase, and superoxide dismutase and 30s ribosomal protein families, **Supplemental Table 8**) were pulled down by pooled total serum IgG from the pre-vaccination timepoint, indicating antibody recognition of these antigens may be relevant to the MOS-specific IgA responses that were associated with natural TB resistance among placebo recipients. Of the remaining 1,078 proteins, a total of 93 antigens detected in MOS were targets of MOS-specific antibodies purified from post-vaccination timepoint sera (**Figure 4A**), ranging from 12-60 proteins per subject. Affinity-purified IgG from individuals who did not develop TB disease enriched a greater number of proteins than sera from definite TB cases (**Figure 4B**), suggesting a narrower response in vaccinated study participants who went on to be diagnosed with TB. Thirty-two of these antigens were targets of post-vaccination serum antibodies in five or more of the 13 participants tested (**Figure 4C**), six of which were annotated as membrane-bound, secreted, or transmembrane proteins in UniProt. Amongst these, ATP synthase, chaperonin GroEL, and UPFO182 are relatively well conserved in Mtb and *M. obuense* (amino acid similarity >79%) (**Supplemental Table 9**). Overall, these data suggest that antibodies to a diversity of MOS proteins were induced by vaccination with polyantigenic whole cell *M. obuense* SRL172.

**Fig. 4:**
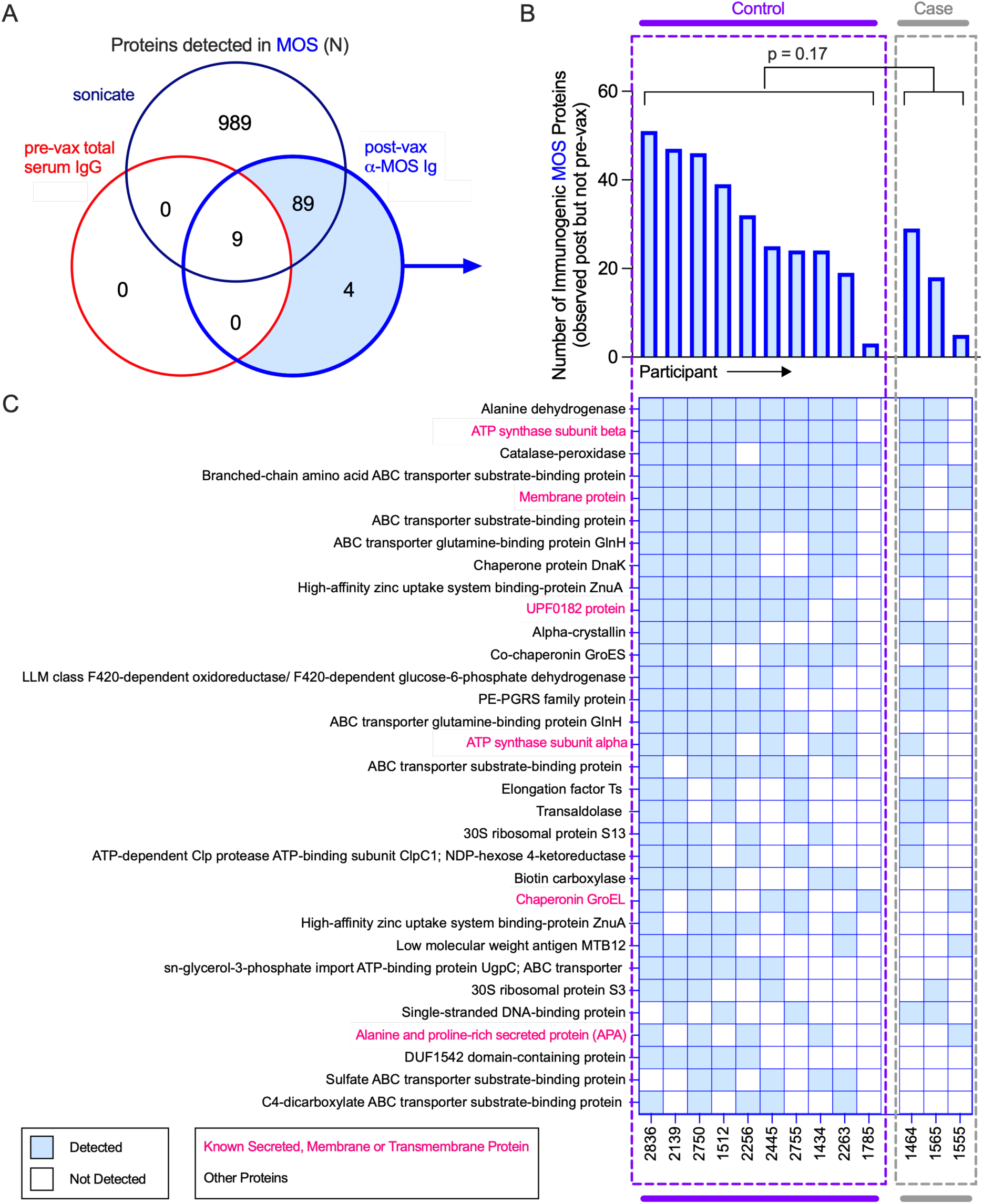
Identification of immunogenic components in MOS. **A.** Venn diagram depicting total proteins detected by mass spectrometry in MOS (navy), or those detected following enrichment with MOS-specific antibodies isolated post-vax (blue) or by pre-vax total serum IgG (red). **B-C.** Total number (**B**) and identity (**C**) of proteins in MOS detected in ≥ 5 of the 10 control and 3 case participants following affinity enrichment using MOS-specific antibody but not pooled IgG by study participant. Statistical significance was determined by Mann-Whitney t test.

### Identification of Mtb proteins recognized by MOS-specific antibodies

We next sought to identify components in Mtb CDC1551 WCL, a well characterized Mtb clinical isolate strain, to which MOS-specific antibodies cross-reacted. MOS-specific antibodies isolated from the serum of nine control and three case study participants with high IgG responses were used to enrich components of Mtb CDC1551 WCL (**Supplemental Figure 10**). Total serum IgG from the pre-vaccination timepoint from seven of these control subjects was employed to identify Mtb proteins recognized by antibodies present before immunization. While more proteins were generally pulled down by post- as compared to pre-immunization sera, similar numbers of proteins were targeted by post-immunization case and control sera, consistent with selection of high responders for this analysis (**Figure 5A**). A total of 1,447 proteins were identified from Mtb CDC1551 WCL, of which 814 were targeted by total serum IgG at pre-vaccination as well as MOS-specific serum antibodies at post-vaccination time points (**Figure 5B**). Relative to the number of immunogenic proteins identified in MOS, this higher number of detected Mtb proteins was surprising, and may be attributable to prior BCG vaccination and Mtb exposure as well as cross-reactive antibodies elicited against common post-translational modifications or heterologous mycobacterial species^51–59^ among other possibilities. Overall, a set of 440 proteins from Mtb CDC1551 WCL were uniquely observed as targets of anti-MOS antibodies at the post-vaccination timepoint (**Figure 5B**).

**Fig. 5:**
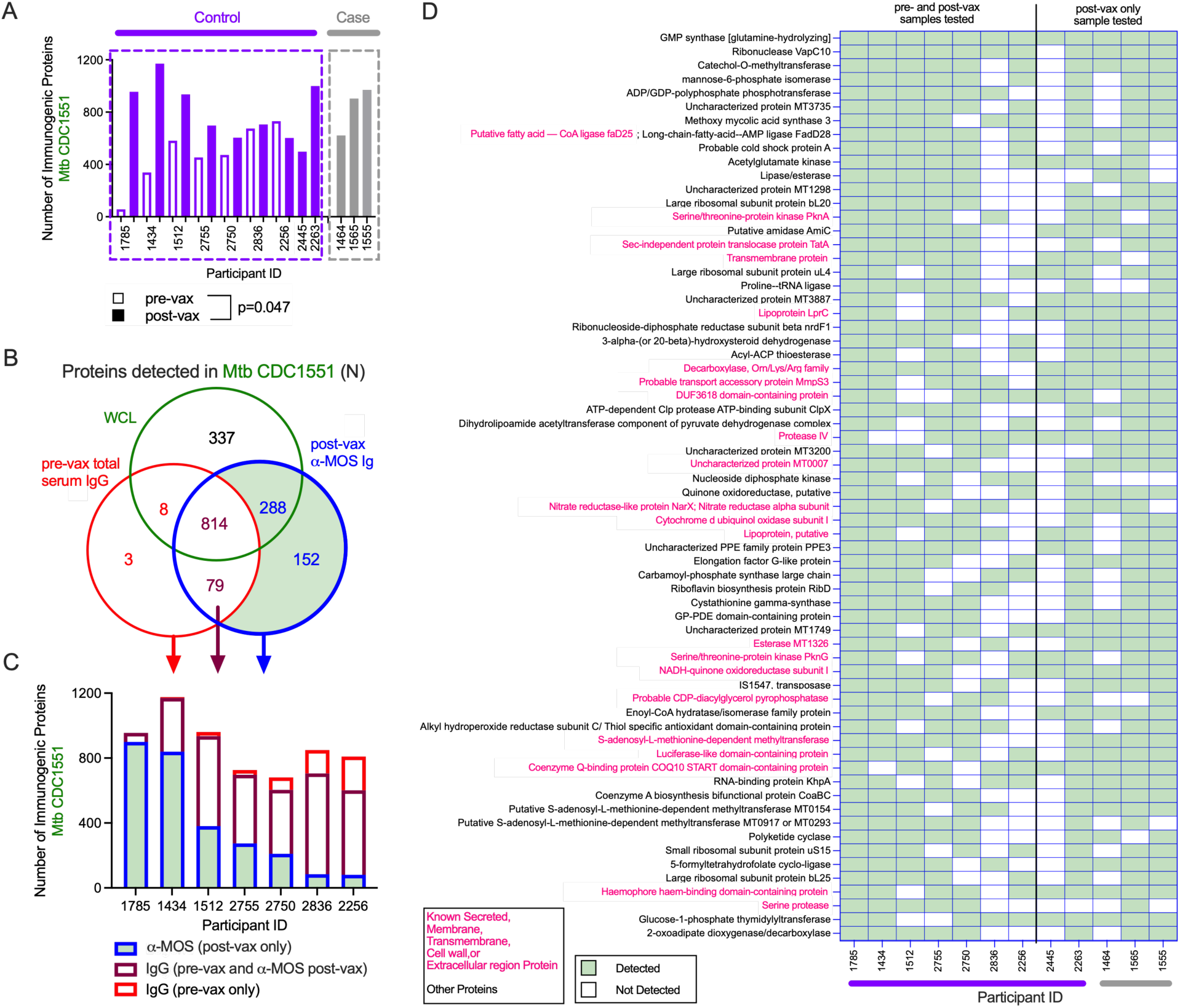
Identification of cross-reactive proteins in Mtb CDC1551 recognized by MOS-specific antibodies. **A.** Number of proteins in Mtb CDC1551 that bound to MOS-specific antibodies isolated from controls (purple) and cases (gray) at post- (filled) vaccination (vax) timepoint and to total serum IgG isolated at pre-vaccination timepoint (hollow). Wilcoxon matched pairs signed rank test was performed to compare number of proteins detected following enrichment by serum antibodies at pre- and post-vaccination timepoint. **B.** Venn diagram depicting the proteins detected in Mtb CDC1551 WCL (green), those detected following enrichment by MOS-specific antibodies isolated post-vax (blue) or by pre-vax serum total IgG (red). **C.** Bar plot depicting the number of Mtb CDC1551 proteins recognized by pre-vax total serum IgG (red), post-vax MOS-specific antibodies (blue), or both (mauve). **D.** Identity of proteins from Mtb CDC1551 detected following enrichment by MOS-specific antibodies post-vax, but not by pre-vax serum IgG in ≥5 participants by participant (controls in purple, n = 8; cases in gray, n = 3).

Per participant, considerable heterogeneity was observed in the number of antigenic protein targets that bound only to MOS-specific antibodies at the post-vaccination timepoint, bound only to total serum IgG isolated at pre-vaccination timepoint, or bound to both (**Figure 5C**). Among the antigenic targets observed only post-vaccination, 66 were observed in five or more of the seven vaccinated control participants (**Figure 5D**), of which 60 met detection criteria in unfractionated Mtb CDC1551 WCL. Of the remaining six, three were observed to be present at low levels but had been filtered out as they were not identified in all three sample injection replicates. Most of these specificities were also enriched by MOS-specific antibodies from the post-immunization timepoint in the two additional controls and three cases for whom pre-immunization samples were not available. Overall, these data suggest that MOS-specific antibodies cross-react with a surprising number of Mtb proteins, consistent with a broad humoral response.

### Quantitative analysis of antibodies to MOS as a CoP

To evaluate how TB disease outcome related to the magnitude of the MOS-specific antibody response, receiver-operator characteristic (ROC) curves, and area under the ROC curve with associated confidence intervals were evaluated for features identified as elevated in controls as compared to cases in both vaccine and placebo groups (**Figure 6A**). MOS-specific IgG1, IgM, as well as the FcγR3A and FcγR2B binding capacity of MOS-specific antibodies in vaccine recipients, and MOS-specific IgA placebo recipients showed significant discriminatory power. In univariate logistic regression models, all tested MOS features presented odds ratios <0.70 among vaccine recipients, and confidence intervals indicated that three features - MOS-specific IgG1 and FcγR3A and FcγR2B ligation showed a statistically significant (q < 0.05) reduction of up to 73% in disease risk per 10-fold increase among vaccine recipients when corrected for multiple hypothesis testing (**Figure 6B**). Similarly, 10-fold greater levels of MOS-specific IgA, the sole CoP among placebo recipients was associated with 79% reduced likelihood of disease (p = 0.014) (**Supplemental Figure 11A**).

**Figure 6.**
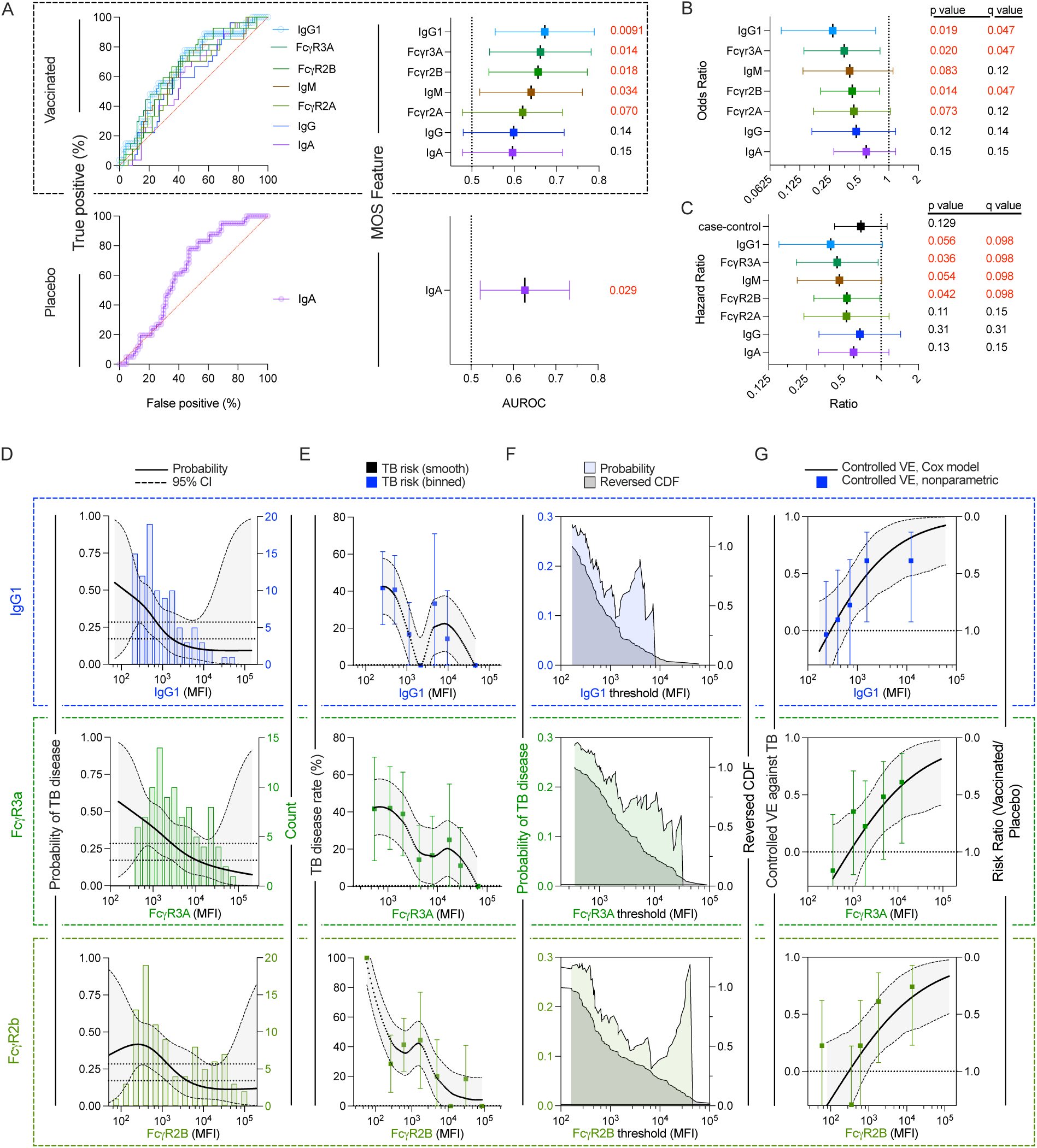
Quantitative analyses of MOS-specific antibodies as correlates of protection. **A**. Receiver-operator characteristic (ROC) curves (left) and areas under ROC (AUROC) (right) for selected MOS-specific antibody markers in vaccine (top) and placebo (bottom) recipients. Error bars indicate 95% Wilson/Brown confidence interval (CI), statistical significance is indicated in inset. **B-C**. Odds ratio (**B**) and Hazard Ratio (**C**) of TB disease per 10-fold increase in each MOS-specific Ab marker. Statistical significance (p value) with Benjamini-Hochberg multiple hypothesis correction (q value) is indicated in inset. Features with values < 0.1 are shown in red. The case-control hazard ratio indicates that observed among the subset of participants selected for case-control dataset, independently of immune markers. **D-G**. Quantitative analysis of MOS-specific IgG1 (top), FcγR3A (center) and FcγR2B (bottom) binding antibodies. **D**. Cumulative incidence of TB disease (probability) by MOS-specific antibody levels, estimated using a marginalized Cox model (black). Dotted lines indicate bootstrap pointwise 95% CI. Graph is overlaid on a histogram of MOS-specific antibody levels (color). **E**. TB disease risk by MOS-specific antibody level smoothed by Loess (gray) or binned (color) with 95% CI indicated. **F**. Cumulative incidence of TB disease (probability) for vaccine recipients with MOS-specific antibody levels above a given threshold value, estimated using nonparametric threshold regression is indicated in color. The estimated cumulative distribution function (CDF) of the marker is indicated in gray. **G**. Controlled vaccine efficacy (VE) as a function of MOS-specific antibody level computed by Cox proportional hazards model (gray) and a nonparametric method assuming that controlled VE is non-decreasing (color). Horizontal dotted line indicates no vaccine efficacy. Error bars or bands indicate 95% CI.

Further, the instantaneous risk of TB disease in vaccine recipients with differing MOS-specific antibody response levels was analyzed by Cox proportional hazard regression. Despite the relatively few participants included in case-control analyses, hazard ratios (HR) for two of these three features also met nominal statistical significance prior to multiple hypothesis correction (**Figure 6C**), indicating that a 10-fold increase the binding of MOS-specific antibody to FcγR2A or FcγR3A was associated with a ∼50% decrease in instantaneous risk of TB disease among vaccine recipients. Supporting the potential validity of these relationships with MOS-specific antibody responses in the context of the limited statistical power of this dataset, while hazard ratio analyses for the entire trial cohort indicated ∼40% efficacy (HR = 0.61, p = 0.03)^9^, an association with only borderline statistical significance was observed among in the small subset of participants included in the case control dataset (HR = 0.69, p = 0.129). Among placebo recipients, MOS-specific IgA presented a statistically significant relationship with TB disease (HR = 0.34, p = 0.023), while again, the case-control participant subset did not (HR = 0.69, p = 0.129) (**Supplemental Figure 11B**). Together, this analysis indicates that MOS-specific antibody responses mark reduced risk of overall and instantaneous TB disease diagnosis.

### TB disease risk decreases and vaccine efficacy increases as MOS-specific antibody marker levels increase

These three MOS-specific antibody features were further analyzed for quantitative relationships with disease risk and vaccine efficacy. Among vaccine recipients, Cox proportional hazard models demonstrated that TB disease probability tended to decrease as the *specific level* of MOS-specific antibody marker level increased, whether probability of disease (**Figure 6D**) or disease rate (**Figure 6E**) was modeled, though sparse data sometimes resulted in large confidence intervals. Furthermore, nonparametric threshold regression analysis estimated the cumulative incidence of diagnosed TB disease for subgroups of vaccine recipients with MOS-specific antibody feature levels *above* a given threshold, which tracked with the reverse cumulative distribution function of MOS-specific antibody levels, suggesting a relatively smooth incremental change in risk with increasing MOS-specific antibodies (**Figure 6F**). Lastly, both Cox and nonparametric models of controlled vaccine efficacy increased with each of these selected MOS-specific antibody features, for example, showing no efficacy at median marker levels observed in the placebo group, to up to a maximum of 92% for the highest observed marker levels (**Figure 6G**). Collectively, this analysis demonstrates the quantitative impact of increasing levels or thresholds of MOS-specific antibody associate with an increasing degree of relative disease resistance.

### Functions of MOS-specific and vaccine-elicited antibodies

Quantitative relationships between biomarkers and outcome are useful for vaccine development and licensure, but markers of efficacy may or may not mechanistically relate to vaccine-mediated protection. To evaluate whether vaccine-induced antibodies might exhibit anti-mycobacterial activity, we conducted *in vitro* tests of antibody effector function on a subset of case and control participant samples. First, we assessed the ability of MOS-specific antibodies to induce human THP-1 monotypes to phagocytose MOS-conjugated fluorescent beads. Post-immunization antibodies showed greater phagocytic activity than those in the same donors at the pre-immunization timepoint, and this activity was greater in vaccinated control than vaccinated case participants, as well as either case or control group placebo recipients (**Figure 7A**). Across all samples tested, as well as within vaccine recipients only, this activity correlated with the level of MOS-specific IgG (**Figure 7B**).

**Figure 7.**
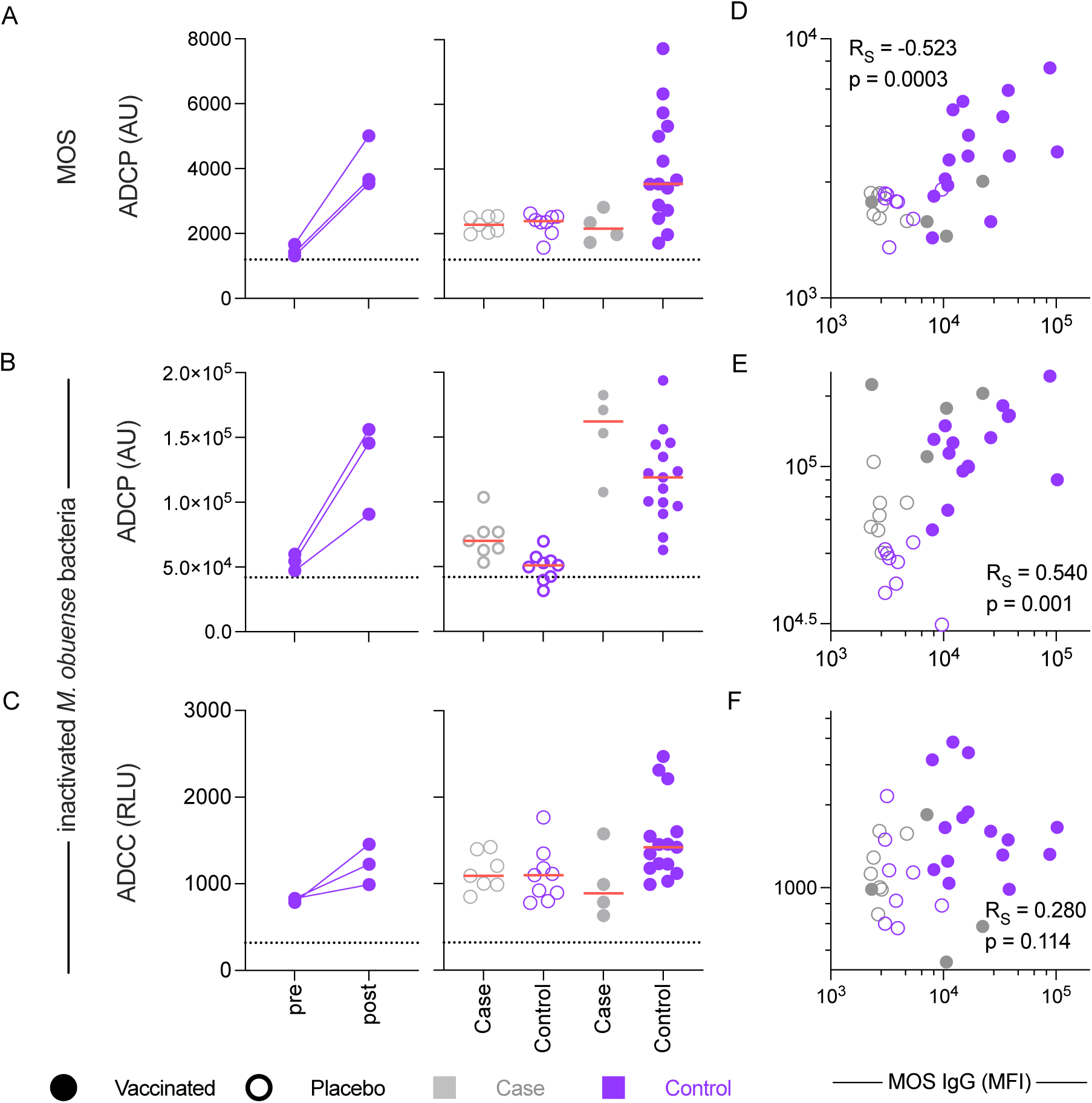
Functions of MOS-specific Abs. **A-C**. Activity in control vaccine recipients before and after immunization (left) and placebo (open symbol) and vaccine (closed symbol) recipient in case (purple, n = 24) and control (gray, n = 11) groups in assays of Antibody-Dependent Cellular Phagocytosis (ADCP) against MOS-conjugated beads (**A**) and inactivated whole *M. obuense* mycobacteria (**B**, MOM, SRL172 vaccine), and Antibody-Dependent Cellular Cytotoxicity (ADCC) as defined by FcγR3A ligation-induced signaling induced by *M. obuense* bacteria (**C**). Dotted horizontal line indicates signal observed in the absence of serum antibody. **D-F**. Correlative relationships between MOS-specific IgG median fluorescent intensity (MFI) and MOS ADCP (**D**), *M. obuense* ADCP (**E**), and *M. obuense* bacteria ADCC (**F**). Spearman correlation coefficient (R_S_) and significance are reported in inset. ADCP activity is given in arbitrary units (AU) and ADCC in relative light units (RLU). Functional testing was performed in biological duplicate. Dotted horizontal line indicates signal observed in the absence of a source of antibody.

To assess activity against mycobacteria, whole inactivated *M. obuense* (SRL172 vaccine particles) were fluorescently labeled, opsonized with antibody, and incubated with phagocytes. In this test as well, serum from vaccine recipients showed greater activity following than before immunization, activity in vaccine recipients exceeded that observed in placebo recipients, and across samples, phagocytosis was correlated with MOS-specific IgG (**Figure 7D,E**).

Lastly, we evaluated FcγR3A signaling induced by antibodies bound to plated mycobacteria as a surrogate of antibody-dependent cellular cytotoxicity (ADCC; **Figure 7F,G**). We observed similar trends between different sample timepoints and donor groups, but no statistically significant correlation with IgG levels and ADCC. Overall, these data suggest that antibodies raised in DarDar vaccine recipients can elicit the effector functions of distinct innate immune receptors and cellular subsets, contributing to interpretation of the levels of MOS-specific antibody responses raised across cohorts evaluated herein (**Supplemental Figure 12**).

## Discussion

Investigational TB vaccines have shown promise to boost or replace childhood BCG immunization^79–81^. Beyond the demonstrated efficacy of vaccination with *M. obuense* SRL172 in the DarDar trial^9^, recent studies of re-routing (i.v.)^35,42^ or re-vaccinating with BCG^5,60^, and novel candidates^61–63^ provide hope for new insights and options to protect vulnerable populations from one of the oldest of human diseases. However, development of effective TB vaccines at all stages is hampered by the lack of established CoP^3,6,61,64^.

We report the novel identification of a humoral CoP by applying systems serology tools to case-control samples from the successful phase III DarDar prevention of disease trial of SRL172 vaccine. Affinity purification and mass spectrometric analysis revealed antibodies to a diversity of immunogenic antigens in MOS and cross-reactivity with those in Mtb. Since epidemiologic studies in humans have demonstrated that infection with non-tuberculous mycobacteria (NTM) can protect against subsequent TB disease, and a number of inactivated whole cell vaccines have been used to prevent TB^52,54,65^, these data suggest a similar mechanism of inducing cross-reactive immunity for inactivated whole cell *M. obuense*.

Not only did IgG, IgM, and IgA antibody responses to the vaccine strain sonicate (MOS) correlate with vaccine-mediated protection from TB disease, among participants who did develop breakthrough TB disease, greater magnitude MOS-specific IgG and IgM responses associated with greater time to disease diagnosis among cases. Generalizability of this human humoral CoP was confirmed by NHP studies of BCG vaccine-mediated protection from TB, and biological relevance suggested by cross-reactivity of MOS-specific antibodies with *M obuense* and Mtb.

In sum, we demonstrate correlations between humoral responses to MOS and protection from TB disease in humans immunized with MOS, humans not immunized with SRL172 (DarDar placebo recipients), as well as BCG-vaccinated NHP. Yet, these correlative data do not identify the mechanism(s) of protection. While antibody effector functions were detected *in vitro*, this study provides no direct evidence that these antibodies or these activities are directly responsible for the protection observed. Tests such as passive antibody transfer or depletion and challenge experiments in animal models would be required to provide evidence that antibodies induced by immunization in the DarDar trial played a mechanistic role in mediating protection from TB disease. Unfortunately, the small sample volumes available and lack of consensus regarding appropriateness of small animal models proved to be practical challenges to pursuing this type of mechanistic evaluation. However, such testing has been conducted in other contexts, resulting in strong mechanistic evidence that i.v. BCG-elicited protection is mediated by T cells in NHP^66^. Increased rates of active disease in both CD4 knock-out mice^67–69^ as well as in humans with low CD4 T cell counts^70,71^ further support the mechanistic importance of cellular responses, which were not evaluated here.

Nonetheless, insight into the targets of MOS-specific antibodies have the potential to refine our understanding of cross-reactivity between NTM and disease-causing mycobacteria in ways that are relevant for TB prevention and treatment. This study did not identify a single or few dominant antigen-specificities that could serve as individual CoP, perhaps due to the polyantigenic nature of whole cell *M. obuense* vaccines and even the possibility that polyantigenic responses to cross-reactive antigens are important to vaccine-mediated protection from TB disease^72^. Our findings do highlight, however, the value that application of proteomic characterization of the antigenicity and immunogenicity of different strains could have to TB vaccine research and development.

Numerous antigenic proteins were identified in both Mtb and *M. obuense* strains. The larger number of Mtb CDC1551 proteins as compared to MOS proteins identified in pull down using MOS-specific antibodies may be attributable in part to the greater characterization of Mtb CDC1551 as compared to *M. obuense* or to differences in immunogenicity between strains, but could alternatively be seen to motivate vaccination with derivatives of this strain. Live-attenuated Mtb vaccines are also being tested, though there are concerns related to safety^73^, and only one such live-attenuated Mtb vaccine (MTBVAC)^74,75^, has advanced in the clinic. Overall, further study to elucidate both targets and mechanism(s) of protection are needed to understand the potential biological relevance of the antigens identified.

We acknowledge limitations of our findings. Study and sample sizes were small, and sufficient volumes of DarDar study samples were not available for all timepoints and tests for the participants selected for case-control analysis. We did not analyze the potentially confounding influence of clinical or demographic variables or stratify humoral responses by the presence or absence of T cell immune responses, so cannot characterize the interplay between the identified humoral CoP and cellular immune responses thought to be critical to protection from tuberculosis. Immune responses observed to relate to disease in the DarDar study also reflect the PLWH population in which the trial was conducted, and in which TB is particularly consequential, but they may or may not generalize to other populations. While we found a clear and consistent pattern of elicitation of MOS-specific antibodies in human and NHP recipients of three TB vaccines, this study cannot assess whether these humoral CoP are associated with protection from TB seen in other vaccines. Such studies are now a high priority next step for TB vaccine development.

Given the exploratory nature of these studies, initial statistical analyses of induced responses did not formally adjust for multiple comparisons but instead relied on comparison of distributions of differences in placebo recipients, or pre-immunization timepoints as study- and dataset-specific means to account for the risk of false discovery. Following the identification of vaccine-induced responses, these corrections were formally applied to support CoP analysis on this more limited panel of features. Among vaccine-induced responses, the robust association of MOS-specific antibody levels with both protection from TB maintained following adjustment argues strongly that the association results from relevance rather than chance. Validation is provided by relationships between the degree of disease risk and vaccine efficacy with MOS-specific antibody levels, as well as by relationships observed with protection mediated by BCG in the NHP model. Potential biological relevance is supported by the observation of cross-reactive antigens between *M. obuense* and Mtb across multiple vaccine platforms and species and the ability of vaccine-induced antibodies to elicit the effector functions of innate immune cell receptors. While we confirmed the elicitation of analogous humoral responses in recipients of the broth-grown whole cell *M. obuense* DAR-901 vaccine, we could not extend CoP analysis to this phase I study since participants were not followed for the development of TB. While overall efficacy against the endpoint of TB infection was not observed in a later study of DAR-901^48^, this does not exclude future study of *M. obuense*-based vaccines given the prior finding of protection from TB disease.

Identification of a humoral CoP for vaccine-induced protection against TB can help accelerate the development of DAR-901 and other promising TB vaccine candidates by providing a new target for immunogenicity assessments and dose finding studies^76^. Further, these findings add to a growing body of evidence motivating reexamination of the role of humoral immunity in marking or contributing to protection from TB disease^24,27,33,77–88^. It will be interesting to extend these studies in other contexts, such as BCG immunization in infants or in BCG re-vac and M72 trials^62,89–91^. Even in the absence of a clear mechanistic underpinning and awaiting further studies of humoral immune responses to other TB vaccine candidates, the identification of an antibody CoP for vaccine-mediated prevention of TB disease in humans may represent a substantial milestone in TB vaccine development.

## Methods

### Serum/Plasma samples

This study evaluated pre- and post-immunization serum samples from participants in the randomized, placebo-controlled, double-blind DarDar phase III^9^ (105 placebo, 95 vaccine recipients, **Supplemental Table 1**) and the randomized, placebo- and BCG-controlled, double-blind, dose ranging DAR-901^47^ phase I (9 placebo, 40 Dar 901, and 9 BCG vaccine recipients, **Supplemental Table 5**) trials. These studies evaluated immunization with either agar- (DarDar) or broth- (DAR-901) grown *M. obuense* in Tanzania and the United States, respectively. Briefly, whereas the DarDar trial included adult residents of Dar es Salaam, Tanzania living with HIV, and who had CD4 cell counts of at least 200 cells/μL and a BCG scar, the DAR-901 study included adult residents of the United States living with or without HIV and with a history of childhood BCG vaccination evidenced by BCG scar. In our case-control sub-study, participants who developed definite TB after immunization were considered as cases and subjects who developed definite TB after immunization were considered as controls. Controls were matched on age, PPD status, and prior TB status. Immunogenicity measures were collected blinded to case-control status. The DarDar study was approved by the Dartmouth Committee for the Protection of Human Subjects, by the Muhimbili University of Health and Allied Sciences (MUHAS) Research Ethics Committee, and by the Division of AIDS Clinical Science Review Committee, National Institutes of Health (NIH). The DAR-901 phase I study was approved by Dartmouth Committee for the Protection of Human Subjects. All participants in both studies gave written informed consent.

Lastly, serum samples from study participants diagnosed with active TB disease (n = 25) or latent TB infection (n = 25) were obtained from Duke University Medical Center (**Supplemental Table 6**). The TB disease group included participants with either culture-confirmed TB (n = 22) or diagnosis per Centers for Disease Control and Prevention criteria (n = 3). Sample collection was approved by the Duke University Institutional Review Board and participants provided written informed consent.

Plasma samples from cynomolgus (*Macaca fascicularis*) or rhesus (*Macaca mulatta*) macaques were collected from studies of BCG immunization^35,42–44^ (**Supplemental Table 7**) performed at University of Pittsburgh, Bioqual Inc., and the National Institutes of Health Vaccine Research Center (VRC). Experimentation and sample collection from each original study was approved by the appropriate local Animal Care and Use Committee, with adherence to guidelines established in the Animal Welfare Act and the Guide for the Care and Use of Laboratory Animals, and the Weatherall Report (8^th^ Edition).

### Fc Array Antibody Profiling

A panel (**Supplemental Table 2**) of recombinant or purified native antigens (Ag 85, APA, α-crystallin, ESAT6, GroES, LAM, Mpt64, and PstS1) as well as complex mixtures (agar-grown *M. obuense* sonicate, broth-grown *M. obuense*, Mtb CDC1551, Mtb 91_0079 whole cell lysates, and cytosolic, membrane, and culture filtrate fractions of Mtb H37Rv) were used for profiling humoral immune responses using a multiplexed binding assay.

Briefly, magnetic carboxylated microspheres (Luminex Magplex) were coupled to each antigen preparation using a two-step carbodiimide reaction as described previously^92^. As needed, preparations were buffer-exchanged into phosphate buffered saline (PBS) using G-25 (Cytivia, 28918004) columns following manufacturer’s protocol to ensure that there were no primary amine groups in the buffer. LAM was modified using DMTMM (4-(4,6-Dimethoxy-1,3,5-triazin-2-yl)-4-methylmorpholinium chloride) (Millipore Sigma, 72104) and coupled using protocols described previously^33,93^.

The levels and Fc characteristics of antigen-specific antibodies were evaluated by multiplex assay as described previously^92^. Serum samples were thawed, diluted in assay buffer (PBS + 1% BSA + 0.1% Tween20), and added to 384 well plates (Greiner bio-one, 781906). A master mixture of antigen-coupled beads was prepared in assay buffer and 45 µL of the mixture was added to each well, such that each well would contain 500 beads per bead region. The final concentration of serum/plasma was 1:100 diluted. The plate was sealed and incubated for 2 hours 15 min at 1,000 rpm at room temperature. Following six washes using an automated magnetic plate washer, bound antigen-specific antibodies were detected with phycoerythrin (PE)-conjugated anti-human or anti-rhesus secondary reagents (**Supplemental Table 10**), or site-specifically biotinylated and tetramerized (Streptavidin-PE (Agilent, Technologies, PJ31S-1)) human Fc receptors (FcγR2A H131, FcγR2B, FcγR3A V158, FcγR3B, FcαR)^94^ at a concentration of 0.65 µg/mL, as described previously^45^. Plates were sealed and incubated for 1 hour 5 min at 1,000 rpm at room temperature, washed 6 times using an automated magnetic plate washer, and beads were resuspended in 50 µL sheath fluid (Luminex™ xMAP Sheath Fluid Plus, 4050021) prior to sealing and agitation at 1,000 rpm for 5 min at room temperature. Data was acquired on a FlexMAP 3D™ (Luminex, Exponent version 4.2.1513), which detected the beads and measured PE fluorescence in order to calculate the median fluorescent intensity (MFI) level for each analyte.

### Immunogenic Peptide Identification

A tandem affinity purification – affinity purification – mass spectrometry (AP-AP-MS) strategy (**Supplemental Figure 10**) was used to first enrich MOS-specific antibodies from post- immunization timepoint serum samples from selected (n=13; 10 control, 3 case) DarDar trial participants that exhibited particularly high MOS-specific antibody responses and for whom sufficient sample was available, and then to capture and identify the MOS components specifically bound by these antibodies.

#### Affinity purification of MOS-specific and control antibodies

Briefly, a 1 mg (i.e., 100 µL) mass of Dynabeads® MyOne™ Carboxylic Acid (Thermo Fisher Scientific, 65012) were conjugated with 50 µg of antigen using two-step carbodiimide reaction. A 100 µL volume of beads was washed with 20 mM MES (2-(N-morpholino)ethanesulfonic acid) (pH 6.0) (Sigma Aldrich, M3671). Post washing, 25 µL of 50 µg/µL of EDC (1-ethyl-3-(3-dimethylaminopropyl)carbodiimide hydrochloride)) (Thermo Fisher Scientific, 22980) and 25 µL of 50 µg/µL sulfo-NHS (N-hydroxysulfosuccinimide) (Thermo Fisher Scientific, 24510), each prepared in ice-cold MES buffer (pH 6.0), were added to the beads, which were then mixed using a vortex and allowed to incubate on a rotational shaker for 30 min at room temperature. The resulting activated beads were washed twice with 150 µL of MES buffer, to which 50 µg of MOS in 50 µL of MES buffer was added. After mixing the sample and the beads, the tube was allowed to incubate on a rotational shaker for 30 min at room temperature. MOS-conjugated beads were then resuspended in 150 µL 50 mM Tris pH 7.4 (Thermo Fisher Scientific, 15567027) for 15 min at room temperature on a rotational shaker. The beads were then washed 4 times with 1X PBS-TBN (Teknova Inc., P0210), resuspended in PBS-TBN, and stored at 4°C. This bead-preparation protocol was scaled according to the quantity of beads needed for processing the desired number of serum samples.

A 100 µL volume of serum sample was mixed with 20 µL of unconjugated beads that had been washed twice with PBS-TBN, and incubated on a rotational shaker for 1 hr 30 min at room temperature. Following depletion of bead-reactive proteins, serum supernatant was withdrawn and then allowed to mix with 20 µL of MOS-conjugated beads at 4°C overnight on a rotational shaker. Unbound serum proteins were removed by washing 3x with PBS-TBN. For elution, the beads were incubated with 50 µL of 1% formic acid (Thermo Fisher Scientific, 28905) on rotational shaker for 10 min at room temperature. The elution step was repeated and eluate fractions combined and protein content estimated by measuring absorbance at 280 nm. Enrichment was confirmed by testing the binding of the eluted anti-MOS antibodies and flow-through serum (various concentrations) to MOS-conjugated Dynabeads® MyOne™ Carboxylic Acid, using flow cytometry, and detection using PE labeled anti-human IgG antibody (Southern Biotech, 9040-09).

As a control, total serum IgG from pre-vaccination time points of vaccinated control participants were isolated by using Melon™ Gel IgG spin purification kit (Thermo Fisher Scientific, 45206) following the manufacturer’s protocol. Due to limited sample availability, total serum IgG could be prepared from only seven of the ten samples from which MOS-specific antibodies were isolated at the post-immunization timepoint.

#### Affinity purification of MOS and Mtb antigens bound by MOS-specific or total serum IgG antibodies

A 25 µg mass of either anti-MOS antibodies (post-vaccination) from individual participants or pooled total serum IgG (pre-vaccination) was conjugated on Dynabeads® MyOne™ Carboxylic Acid as described above. To identify the immunogenic components of MOS, antibody-conjugated beads were incubated with 50 µg MOS in 50 µL PBS-TBN overnight on a rotational shaker at 4°C. The beads were then washed and the components in MOS that bound to the anti-MOS antibodies were subsequently eluted using 1% formic acid and 0.5M sodium phosphate dibasic for analysis by mass spectrometry.

To identify the components of Mtb that were recognized by MOS-specific antibodies, Dynabeads® MyOne™ Carboxylic Acid beads conjugated with MOS-specific antibodies (post-vaccination), or, as a control, total IgG from serum (pre-vaccination) from individual participants, were incubated with Mtb CDC1551 WCL (BEI NR-14823) (50 µg in 50 µL PBS-TBN). Similar steps as described above were followed to affinity purify and elute antigens in Mtb WCL that bound to MOS-specific antibodies.

#### Mass spectrometric (MS) sample preparation

Eluted components, or as controls, 20 µg total MOS or Mtb CDC1551 (BEI NR-14823), to be analyzed on mass spectrometer were mixed with MS grade water (Thermo Fisher Scientific, 51140) to make a final volume of 65 µL. To this solution, 65 µL of 100% TFE (2,2,2-trifluoroethanol) (Honeywell, 0584150ML) and 6.5 µL of 100 mM DTT (dithiothreitol) (Roche, 10708984001) was added, followed by heating at 55 °C for 45 min. After this denaturation step, the sample was allowed to cool down at room temperature for 15 min prior to addition of 3.9 µL of 550 mM IAM (iodoacetamide) (VWR, M216-30G) and incubation in the dark at room temperature for 30 min. Samples were then diluted with 1159.6 µL of 40 mM Tris hydrochloride (pH 7.5) and trypsin (1:30 (w/w) trypsin/protein) (ThermoFisher Scientific, 90059) was added prior to incubation at 37 °C for approximately 14 hours. The trypsin digestion was stopped by adding 13 µL of 100% formic acid to the solution. Sample volume was reduced under vacuum in a centrifuge concentrator (Eppendorf, 022820168) so that the final volume was approximately 150 µL. Sample cleanup was carried out using Pierce™ C18 Spin tips (Thermo Fisher Scientific, 84850). The peptides were allowed to bind to tips, washed three times with 0.1% formic acid, eluted in a buffer containing 60% acetonitrile and 40% of 0.1% formic acid solution in low protein binding tubes (ThermoFisher Scientific, 90410), and allowed to dry under vacuum. N-linked glycans were cleaved by addition of 500 units of glycerol-free PNGase F (New England Biolabs, P0709S) per 20 µg of protein and the total sample volume was adjusted to 17 µL using 100 mM ammonium bicarbonate (Millipore Sigma, A6141) for incubation at 37 °C for 1 hour. A 2 µL volume of 1% formic acid and 1 µL acetonitrile (ThermoFisher Scientific, 85188) was added, and samples were transferred into mass spectrometry injection vials (ThermoFisher Scientific, 6ERV1103PPC and 6ARC11ST10R). This protocol was scaled according to the concentration of protein in the sample as measured by absorbance at 280 nm.

Samples were analyzed by liquid chromatography-tandem mass spectrometry using an Easy-nLC 1200 (ThermoFisher Scientific) connected to an Orbitrap Fusion Tribrid (ThermoFisher Scientific, Orbitrap Fusion Tune Application 4.2.4321). Peptides were passed through a PepMax RSLC C18 (ThermoFisher Scientific, 164946) prior to separation on an EasySpray HPLC column (ThermoFisher Scientific, ES903) using a 1.6%–76% (v/v) acetonitrile gradient over 90 mins at 300 nL/min. Eluted peptides were injected into the mass spectrometer using an EASY-Spray source. Peptides were analyzed in data-dependent mode with parent ion scans collected at a resolution of 120,000in the orbitrap. Monoisotopic precursor selection and charge state screening were used. Ions with charges ≥+2 were selected for collision-induced dissociation (CID) fragmentation, and MS2 spectra were acquired in the ion trap, with a maximum of 20 MS2 scans per MS1. A dynamic exclusion duration of 15-s was used to exclude ions selected more than twice in a 30-s window. Each sample was injected three times to generate technical replicate datasets.

#### MS – MS data analysis

Protein sequence databases were constructed by downloading the *M. obuense* and Mtb CDC1551 proteomes from UniProt^95^. The sequence database of each organism was merged with a list of common protein contaminants (MaxQuant) to make the final database, against which the spectra was searched using SEQUEST (Proteome Discoverer 2.4, ThermoFisher Scientific). Searches considered fully tryptic peptides only and allowed up to two missed cleavages. A precursor mass tolerance of 5 ppm and fragment mass tolerance of 0.5 Da were used. Modifications of carbamidomethyl cysteine (static) and oxidized methionine, and formylated lysine, serine, and threonine (dynamic) were selected. High-confidence peptide-spectrum matches (PSMs) were filtered at a false discovery rate of <1% as calculated by Percolator (q-value <0.01, Proteome Discoverer 2.4; ThermoFisher Scientific). For each scan, PSMs were ranked first by posterior error probability (PEP), then q-value, and finally XCorr. The average mass deviation (AMD) for each peptide was calculated as described previously^96^, and peptides with

|AMD| >1.5 ppm were removed. Ambiguous peptide mass matches (i.e. peptides that matched with <90% amino acid identity to more than one unique protein without homologous regions) were removed. For each peptide, a total extracted-ion chromatogram (XIC) area was calculated as the sum of all unique peptide XIC (extracted ion chromatogram) areas of associated precursor ions. The detected peptides were filtered such that the PSM count was ≥3. A protein was considered to be detected if at least one of its fragments was detected during mass spectrometry. A list of proteins detected in samples was made. The lists of proteins detected in pre- and post-immunization timepoints were compared in order to identify those that were detected at the post-vaccination timepoint but were not observed at the pre-vaccination timepoint.

### Antibody functions

#### Antibody mediated cellular phagocytosis (ADCP)

Antibody-dependent phagocytic activity was determined by quantification of antigen-conjugated fluorescent beads by the monocytic human THP-1 cell line^97^. Fluorescent polystyrene 1.0 µm beads (Life Technologies) were conjugated with MOS via crosslinking with primary amines. Following washing, 10^5^ beads per well were incubated in media [RPMI1640 (Fisher) + 10% FBS (Biowest] with participant serum samples at 1:100 dilution in media. THP-1 cells (ATCC) at a 1:10 cell to beads ratio and incubated at 37°C under 5% CO_2_ for 4 hours. Cells were washed in cold PBS, fixed with 4% paraformaldehyde, washed with PBS, and analysed on a flow cytometer (Advanteon Novocyte, NovoExpress 1.6.2). Analysis included gating on the cell population in the FSC vs SSC plot, followed by identification of the proportion of cells that were bead+ and the median fluorescent intensities (MFI) of this population. A phagocytosis score was calculated by multiplying the MFI value by the percent of bead+ cells.

For the whole inactivated *M. obusense* mycobacteria (MOM, SRL-172 vaccine particles), sulfo-NHS-LC-biotin (Thermo Scientific) was tetramerized with streptavidin-RPE (Invitrogen) at a 4:1 molar ratio in ultrapure water by mixing end-over-end for 20 minutes at room temperature. MOM was then conjugated to this sulfo-NHS-biotin-streptavidin-RPE fluorescent marker via crosslinking with primary amines on the surface of the mycobacteria. Particles were washed via centrifugation and resuspended in PBS. Following washing, labelling was confirmed and vaccine particles were counted using a flow cytometer (Advanteon Novocyte, NovoExpress 1.6.2). To test antibody phagocytic activity, 10^5^ fluorescently-labelled vaccine particles were added to each assay well and the assay was performed as described above. Assays were conducted in biological and technical duplicate.

#### Antibody-dependent cellular cytotoxicity (ADCC)

An FcγR3A stimulation reported cell assay was used to model ADCC activity. Assay plates were prepared by coating high-binding flat bottom 96-well plates (VWR) with 50 µL per well of whole inactivated *M. obusense* mycobacteria (SRL-172 vaccine particles) in PBS and incubated overnight at 4°C. Assay plates were washed with PBS and then blocked using PBS + 2.5% bovine serum albumin (Sigma) for 1 hour at room temperature. Assay plates were washed with PBS and 20 µL of participant serum, diluted at 1:100 in PBS, were added to each well. A 20 µL volume of PBS was used as a negative control. Assay plates with serum were allowed to incubate for 20 min at 37°C. Jurkat-Lucia NFAT-CD16 cells (Invivogen) were cultured in growth medium - IMDM media supplemented with 10% fetal bovine serum, 1 mM sodium pyruvate, 1x non-essential amino acids, 100 µg/mL Normocin, 100 µg/mL Zeocin, 10 µg/mL Blasticidin. Cells were pelleted by centrifugation at 300 x g for 7 minutes and resuspended in IMDM + 10% fetal bovine serum, 1 mM sodium pyruvate, and 1x non-essential amino acids. A 180 µL volume containing 100,000 cells was added to each well. Assay plates were incubated for 24 hours at 37°C with 5% CO_2_. Following incubation, 25 µL from each well was mixed with 75 µL of Quanti-Luc luciferase substrate (Invivogen) and luciferase signal was measured on a plate reader (SpectraMax, Molecular Devices, Softmaxpro7.0.2). Assays were conducted in biological and technical duplicate.

### Analysis of Biomarker Relationships to TB Disease Outcome

#### Statistical Methods

MOS-specific antibody levels (IgG, IgG1, IgA, IgM, FcγR2a, FcγR2b, FcγR3a) were log₁₀-transformed prior to all analyses. Analyses were restricted to participants with complete antibody data at the post-immunization timepoint and valid follow-up. The analytical approach was adapted from previously described correlates of risk frameworks^90,98–100^.

#### Discrimination

Receiver-operating characteristic (ROC) curves and area under the ROC curve (AUROC) were estimated separately for vaccinated and placebo recipients to assess the discriminative ability of each antibody feature. AUROCs are reported with 95% Wilson/Brown confidence intervals and two-sided p-values for confidence intervals in GraphPad Prism.

#### Odds and Hazard Ratio Calculation

The association between each antibody marker and TB infection was assessed using two complementary approaches. First, logistic regression was used to estimate odds ratios (OR) per 10-fold increase in antibody titer, with predicted TB disease probability curves derived from natural cubic spline fits. Second, Cox proportional hazards regression was used to estimate hazard ratios (HR) per 10-fold increase, with follow-up time defined as days from the MOS visit to TB diagnosis or censoring. Both models were fitted on the log₁₀-transformed marker scale. Multiple comparisons across the seven analytes were addressed using the Benjamini-Hochberg false discovery rate procedure; q < 0.05 was considered statistically significant.

#### Risk Visualization

Three complementary visualizations were produced for each analyte. First, a histogram of antibody levels was overlaid with logistic and Cox-derived probability curves to show the distribution of responses alongside infection risk across the full marker range. Second, a threshold analysis panel showed the empirical infection probability and reversed cumulative distribution function (CDF) as a function of each candidate threshold, with Cox-derived hazard ratios superimposed on a secondary axis. Third, a risk stratification panel divided the marker range into eight equal-width bins and overlaid a loess-smoothed risk curve on the Cox-derived cumulative incidence curve to compare model-based and empirical risk estimates.

#### Controlled Vaccine Efficacy

Controlled vaccine efficacy (VE) was estimated as a function of antibody level, defined as the proportional reduction in TB risk for a vaccinated individual relative to the placebo arm incidence rate. CVE was estimated using both a Cox proportional hazards model and a nonparametric approach that constrains efficacy to be non-decreasing with increasing antibody level, following Gilbert et al.^101^ and Kenny et al.^98^.

### Software

Data was analyzed and graphed using Graph Pad Prism (version 9.4.1 and version 10.2.1) and Rstudio (version 4.2.1 and 4.2.3) ggplot2^102^, tidyverse^103^, dpylr^104^, ggpubr^105^ and ggVennDiagram^106^. Correlate analyses were implemented in R (v4.1.0+) using the *survival*, *splines*^107^, and *pROC* packages. ROC analyses were additionally performed in GraphPad Prism using the Wilson/Brown method for confidence intervals. Mass spectrometry data was analyzed using SEQUEST and Percolator (Proteome Discoverer 2.4). Statistical tests (two-sided unless otherwise noted) and multiple hypothesis corrections performed are defined in figure and table legends.

## Data Availability

The raw proteomic data has been deposited in MassIVE (https://massive.ucsd.edu/ProteoSAFe/static/massive.jsp) under accession ID MSV000094946. Source data are provided are provided with this paper: the raw data for measurement of immune responses from human subjects and from NHP studies are available in **Data Files 1-4**. Data graphed in figures are available in SOURCE DATA File 1.

## Code Availability

Code used in analysis is available from prior work^90,98–107^, as noted in greater detail above.

## Acknowledgements

We would like to acknowledge the DarDar and DAR-901 study teams and participants for their contributions, as well as BEI Resources, NIAID, NIH for providing Mtb antigens. The anti-rhesus IgA (9B9) was provided by the NIH Nonhuman Primate Reagent Resource (NIAID U24 AI126683).

## Author Contributions

N.S.K performed the systems serology experiments and analyzed the corresponding data. N.S.K. and N.C.C. performed the mass spectroscopy experiments and analyzed the corresponding data.

U.S.K. performed function antibody tests. O.G.P. performed statistical analysis. T.P.L., W.W.A and C F.v.R provided serum samples and clinical data for DarDar and DAR-901 trial participants.

J.E.S. provided serum samples and clinical data for subjects diagnosed with TB disease and TB infection. P.A.D, R.A.S. and M.R. provided rhesus macaque plasma samples and metadata. E.C.L., S.J. and C.A.S. provided cynomolgus macaque plasma samples and metadata. J.L. provided guidance to analyze the mass spectrometric data. J.E.S., T.P.L, P.A.D., R.A.S, M.R. and C.F.v.R provided domain guidance in the field of tuberculosis during this study. T.P.L., F.v.R, N.S.K and M.E.A. designed the DarDar case-control sub-study. N.S.K. and M.E.A. designed the experiments. N.S.K. prepared the figures and drafted the manuscript. M.E.A. and J.L. supervised the research. N.S.K. and M.E.A. finalized the manuscript. All authors reviewed and edited the manuscript.

