## Supplemental Materials for "Humoral correlate of vaccine-mediated protection from tuberculosis identified in humans and nonhuman primates"

| **Supplemental Figures and Legends** | |
|---|---|
| Supplemental Figure 1 | Antibody features that differ in SRL172 vaccine recipients as compared to placebo controls by timepoint |
| Supplemental Figure 2 | Differential immune responses in vaccinated and placebo groups in DarDar trial |
| Supplemental Figure 3 | CD4 cells and HIV-1 viral load in DarDar trial participants at pre-vaccination timepoint |
| Supplemental Figure 4 | Longitudinal profile of MOS-specific Correlates of Protection in vaccinated and placebo recipients in the DarDar trial |
| Supplemental Figure 5 | TB acquisition over time in high and low MOS-specific antibody responders in breakthrough Placebo Cases |
| Supplemental Figure 6 | MOS specific IgG responses in participants from DarDar and DAR-901 trial |
| Supplemental Figure 7 | MOS-specific IgM and IgA responses in BCG immunized non-human primates |
| Supplemental Figure 8 | MOS-specific IgG response by route and dose |
| Supplemental Figure 9 | Correlation between MOS-specific IgG and disease outcomes |
| Supplemental Figure 10 | Schematic for immunoproteomics |
| Supplemental Figure 11 | Placebo group correlate analysis |
| Supplemental Figure 12 | Summary of cohorts used and key observations |
| **Supplemental Tables** | |
| Supplemental Table 1 | Characteristics of case control cohort of subjects from DarDar trial |
| Supplemental Table 2 | Antigens used for Fc Array Assay |
| Supplemental Table 3 | Identification of vaccine-induced antibody responses |

| | |
|---|---|
| Supplemental Table 4 | Case-control analysis of vaccine-induced immune responses in the vaccinated group at post-vaccination timepoint |
| Supplemental Table 5 | DAR-901 trial study cohort |
| Supplemental Table 6 | TB disease and TB infection study cohort |
| Supplemental Table 7 | Non-human Primate cohorts |
| Supplemental Table 8 | MOS proteins pulled down by pre-immunization serum antibodies |
| Supplemental Table 9 | Sequence similarity between select *M. obuense* proteins and Mtb proteins |
| Supplemental Table 10 | Fc Array detection reagents |
| **Data files** | |
| Data File 1 | Immune responses measured in DarDar trial participants |
| Data File 2 | Immune responses measured in DAR-901 trial participants |
| Data File 3 | Immune responses measured in active and latent TB cohort |
| Data File 4 | Immune responses measured in non-human primate studies |

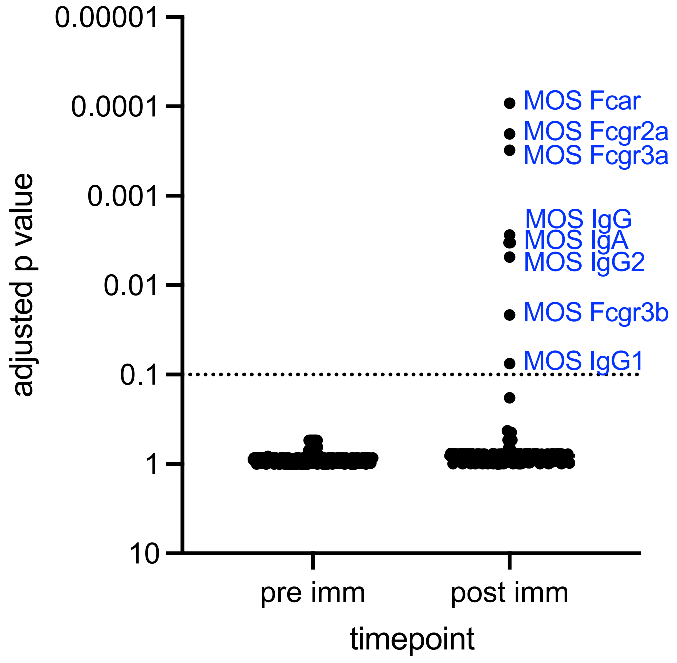

**Supplemental Figure 1: Antibody features that differ in SRL172 vaccine recipients as compared to placebo controls by timepoint.** Features induced in SRL172 recipients, as defined as those that exhibited p<0.10 when analyzed in the context of multiple hypothesis correction (False Discovery Rate with Benjamini Hochberg method) at pre-vaccination (left) and post-vaccination (right) timepoints between placebo and SRL172 immunized participants. Dotted line indicates FDR = 0.1.

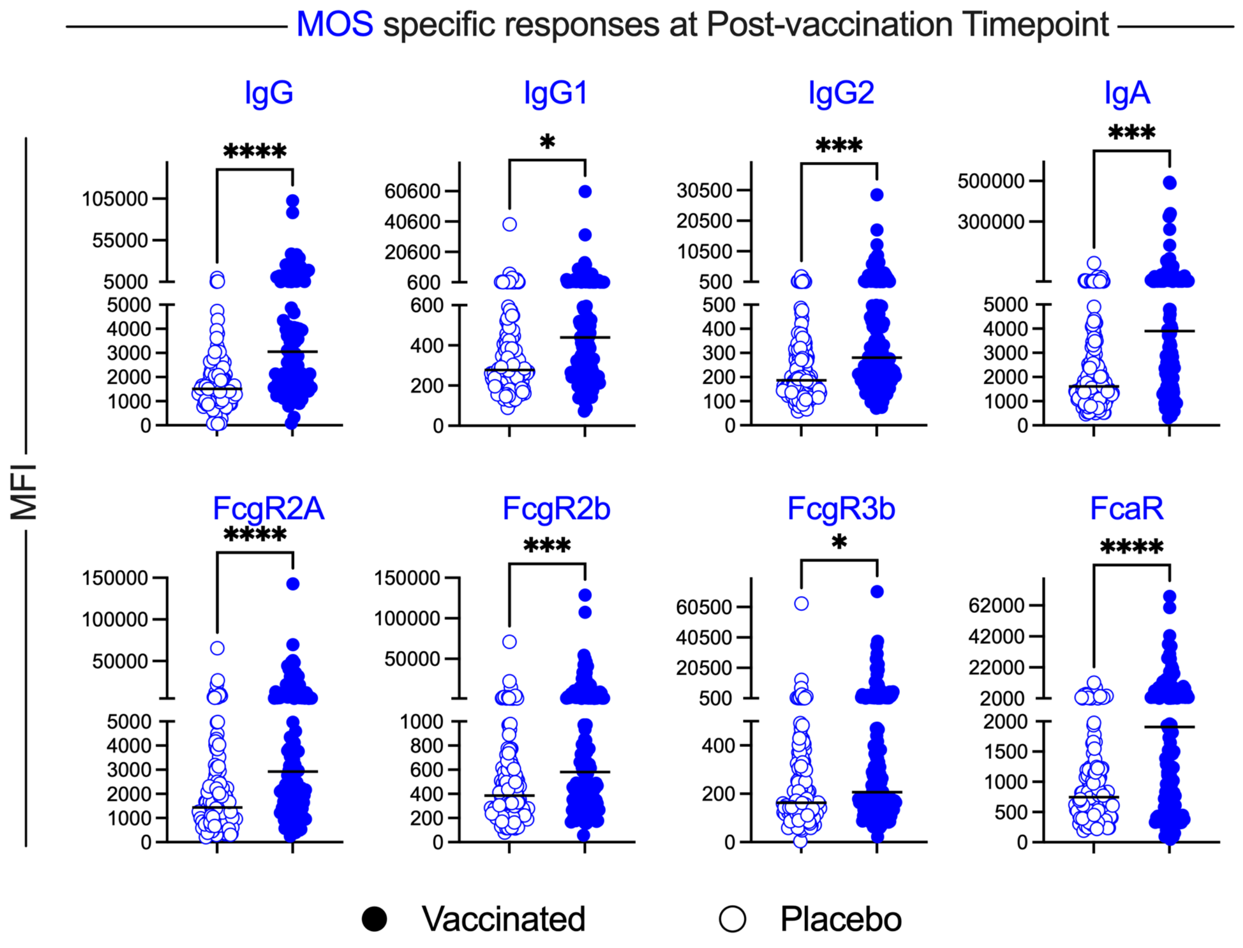

**Supplemental Figure 2: Differential immune responses in vaccinated and placebo groups in DarDar trial.** Box plots comparing the levels of select immune features at post-vaccination timepoint amongst placebo (hollow circles) and vaccinated (solid circles) participants. Statistical significance was determined by Welch's t-test (****p<0.0001, ***p<0.001, **p<0.01, *p<0.05, ns p ≥ 0.05). Bar indicates median.

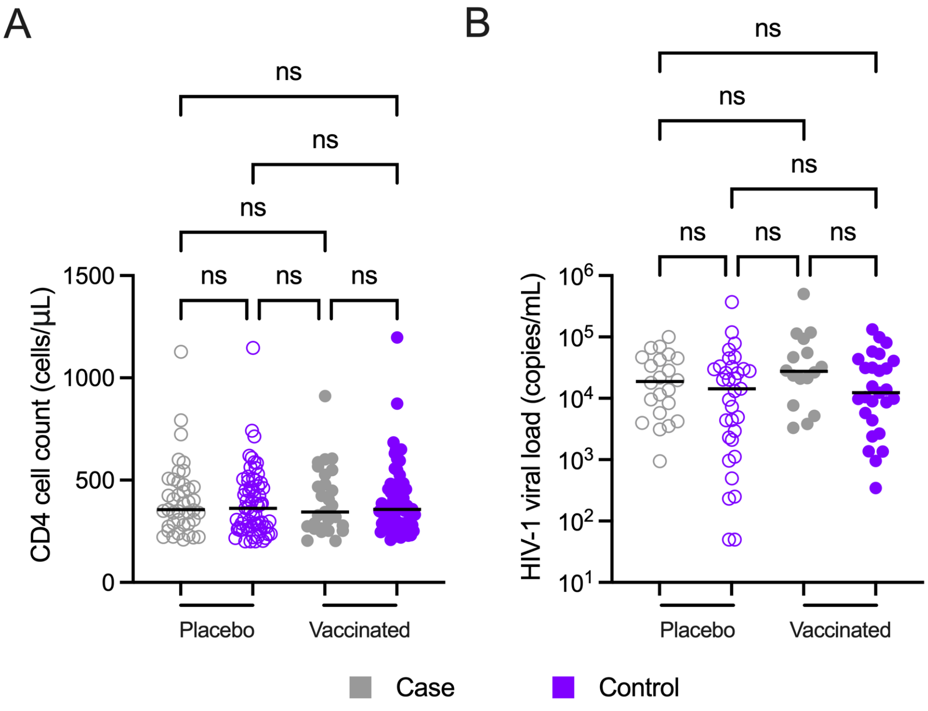

**Supplemental Figure 3: CD4 cells and HIV-1 viral load in DarDar trial participants at pre-vaccination timepoint.** CD4 T cells and HIV-1 viral load in cases (gray) and controls (purple) in vaccinated (solid circles) and placebo (hollow circles) groups at pre-vaccination timepoint. Statistical analysis was calculated by Kruskal-Wallis test (ns $p \geq 0.05$). Bar indicates median.

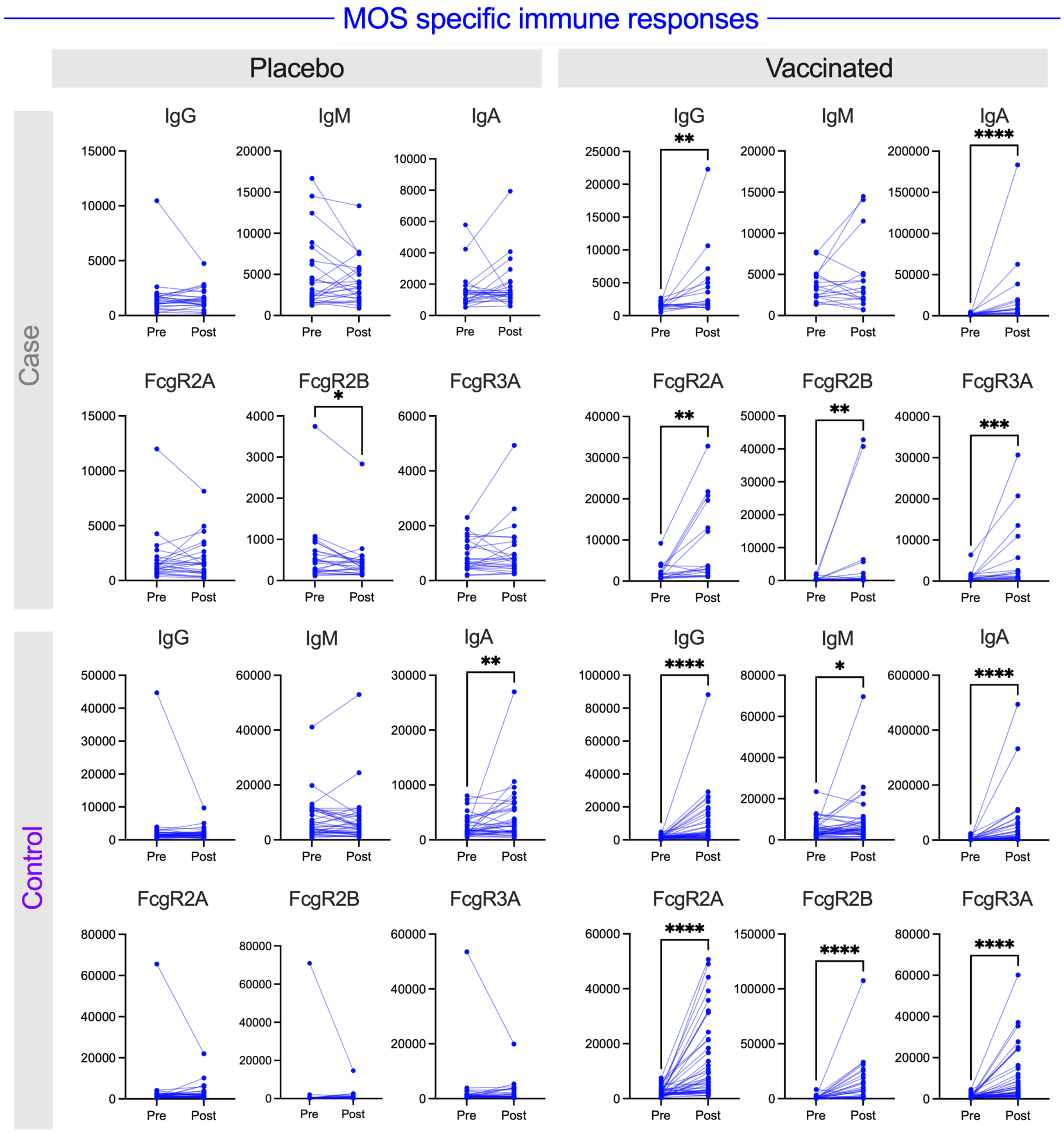

**Supplemental Figure 4: Longitudinal profile of MOS-specific Correlates of Protection in vaccinated and placebo recipients in the DarDar trial.** Longitudinal profiling of select MOS-specific immune features identified as correlates of Protection by Case-control analysis in placebo (left) and vaccinated groups (right), amongst cases (top) and controls (bottom). Statistical analysis was performed by Wilcoxon matched paired sign rank test (****p<0.0001, ***p<0.001, **p<0.01, *p<0.05).

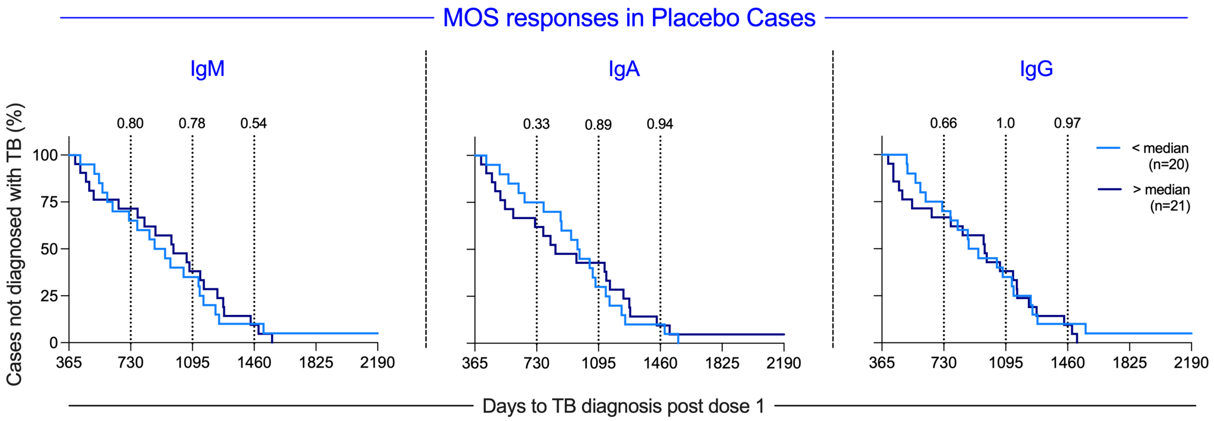

**Supplemental Figure 5: TB acquisition over time in high and low MOS-specific antibody responders in breakthrough Placebo Cases.** Kaplan-Meier curves depicting diagnosis of TB over time in placebo recipient cases for high (≥ median, dark blue) and low (≤ median, light blue) MOS-specific IgM (left), IgA (center) and IgG (right) responders. Statistical significance was evaluated at years 1, 2, and 3 (vertical lines) post final vaccine dose using log-rank (Mantel-Cox) test.

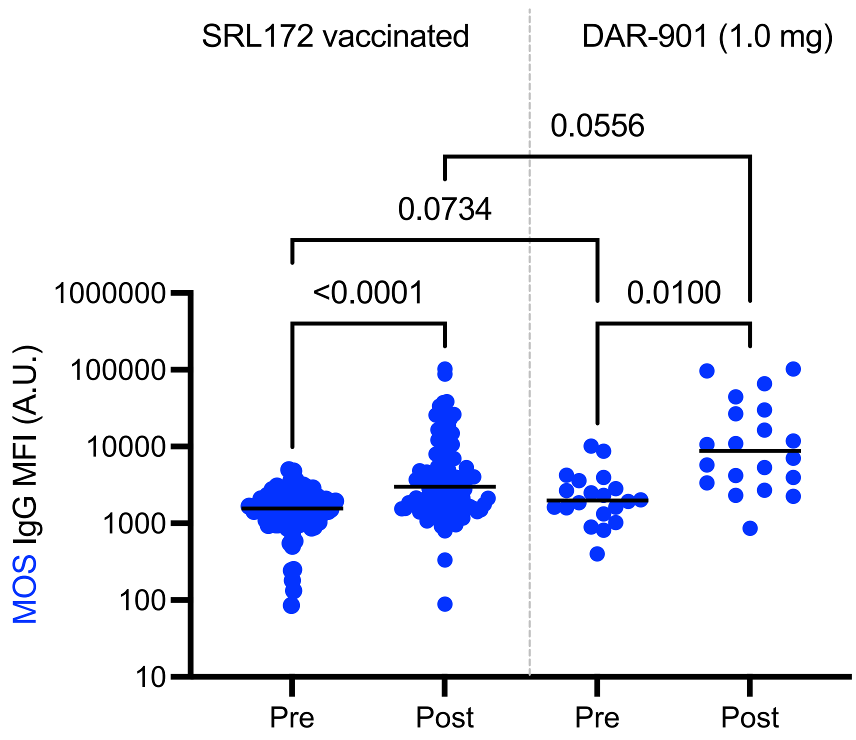

**Supplemental Figure 6: MOS specific IgG responses in participants from DarDar and DAR-901 trial.** MOS specific IgG responses vaccinated participants of the DarDar trial who received SRL172 vaccine and subjects administered with 1.0 mg DAR-901 vaccine at pre- and post- vaccination timepoint. Statistical significance was determined by using ANOVA.

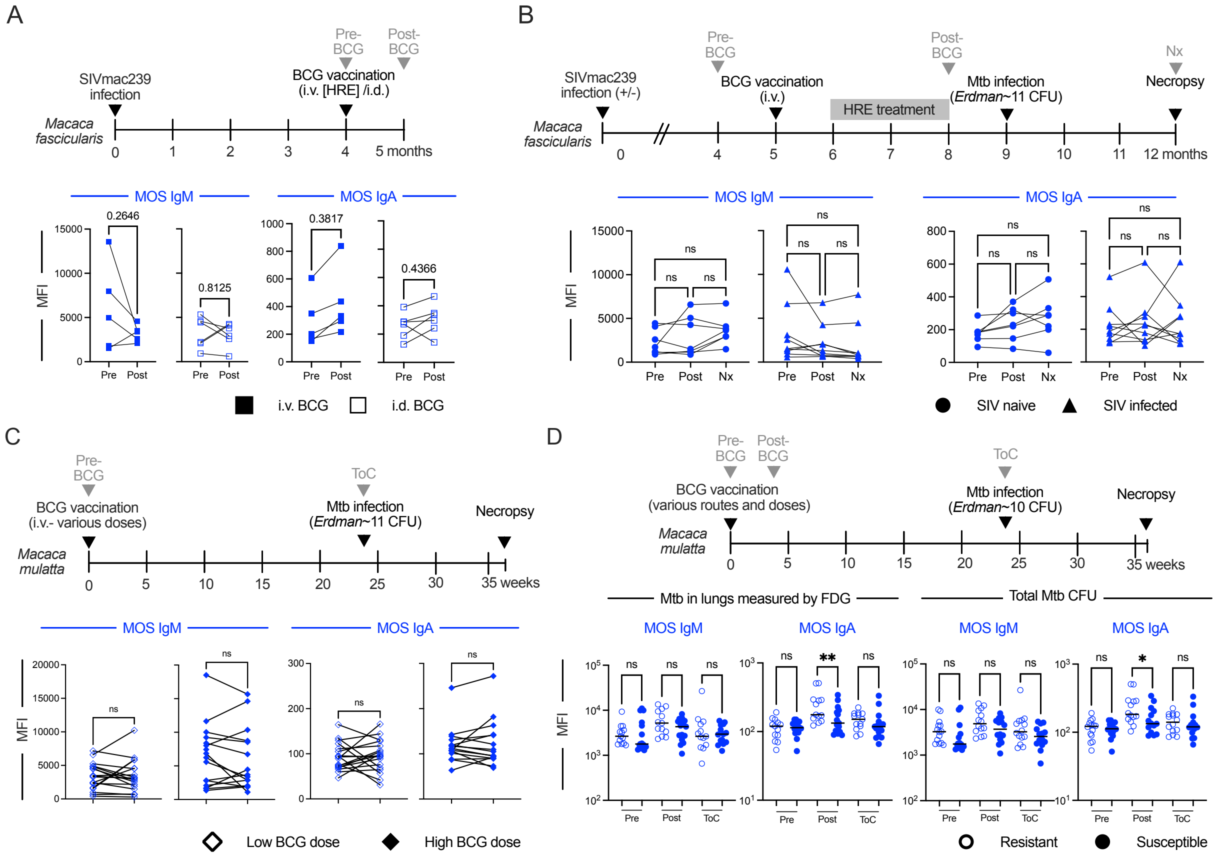

**Supplemental Figure 7: MOS-specific IgM and IgA responses in BCG immunized non-human primates. A.** Longitudinal profiling at pre- and post-vaccination timepoints of MOS specific IgM and IgA responses in SIV-infected *M. fascicularis* administered i.d. or i.v. BCG. Statistical significance was determined by paired t-test. **B.** Longitudinal profiling at pre-vaccination, post-vaccination and necropsy (Nx) timepoints of MOS-specific IgM and IgA responses in SIV-infected and naïve *M. fascicularis* who were administered i.v. BCG and protected from Mtb challenge. Statistical significance was determined by paired one-way ANOVA. **C.** Longitudinal profiling of MOS-specific IgM and IgA responses at pre-vaccination and time of Mtb challenge (ToC) in *M. mulatta* vaccinated with low and high dose i.v. BCG. Statistical significance was determined by paired t-test. **D.** Longitudinal profiling of MOS-specific IgM and IgA responses in *M. mulatta* by Mtb infection burden determined by FDG imaging (left) and by colony forming units of Mtb in lungs (right). Statistical significance was determined by one-way ANOVA. Significance is indicated as: $^{**}p<0.01$, $^{*}p<0.05$, ns $p\geq0.05$.

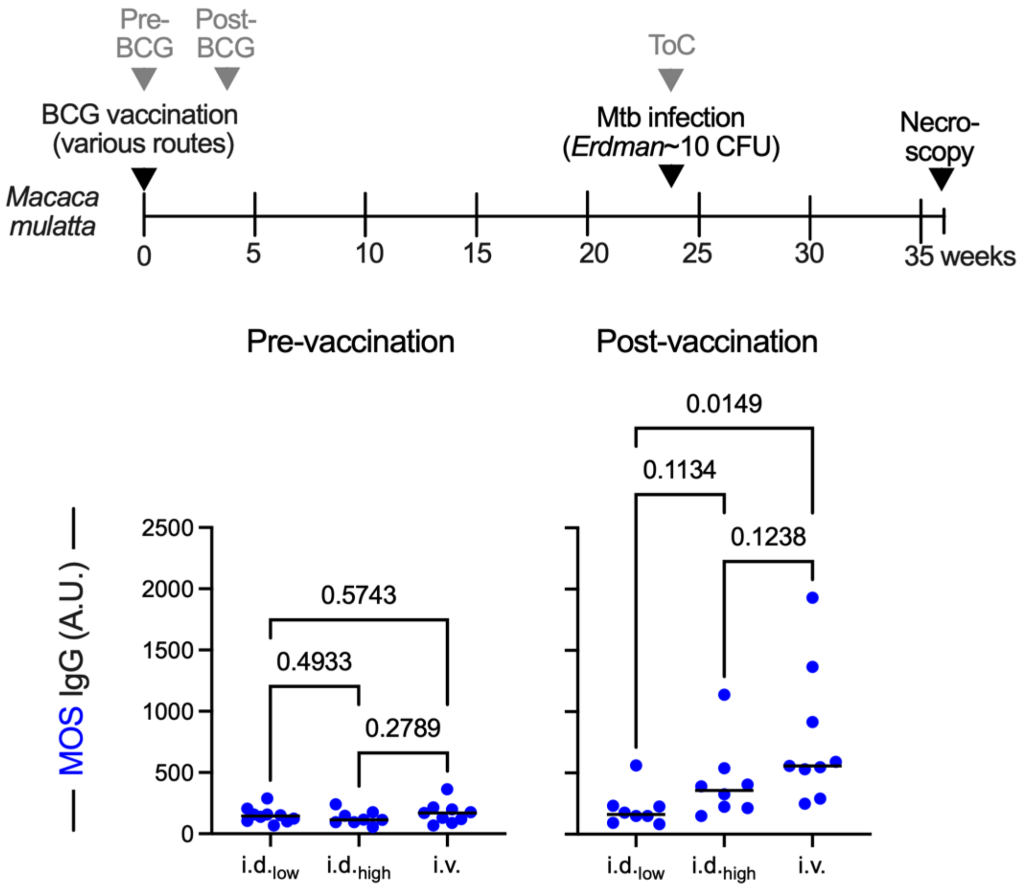

**Supplemental Figure 8: MOS-specific IgG response by route and dose. A.** Profiling of MOS-specific IgG responses in *M. mulatta* vaccinated with different routes (i.d. and i.v.) and dose of BCG vaccination. Statistical significance was determined by one-way ANOVA. Significance is indicated as: **p<0.01, *p<0.05, ns p≥0.05.

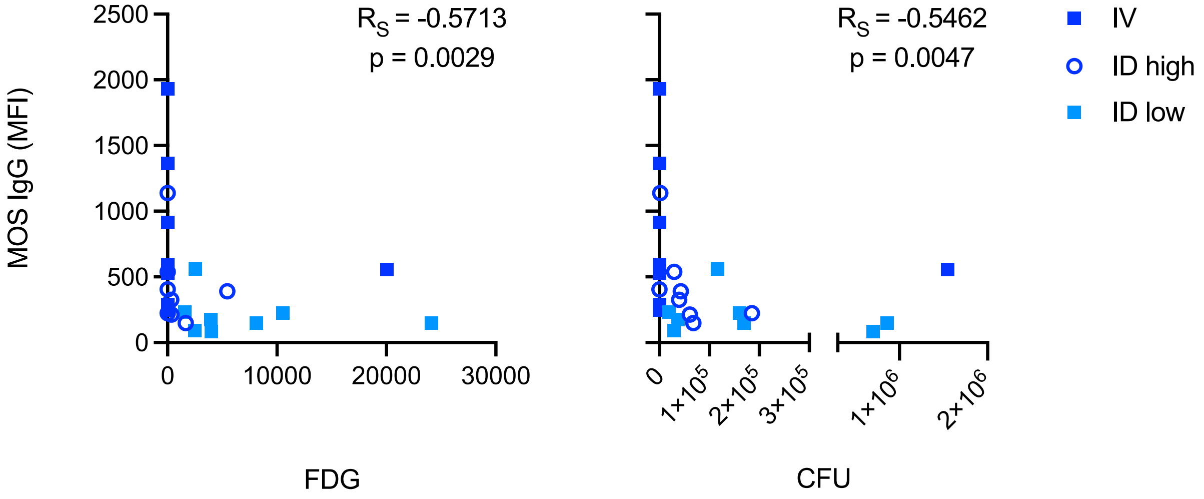

| | FDG vs. ID high | FDG vs. ID low | FDG vs. IV |
|---|---|---|---|
| $R_S$ | -0.5834 | -0.2635 | 0.000 |
| p | 0.1369 | 0.5240 | >0.9999 |
| | ns | ns | ns |
| | Exact | Exact | Exact |

| | CFU vs. ID high | CFU vs. ID low | CFU vs. IV |
|---|---|---|---|
| $R_S$ | -0.8333 | -0.4311 | 0.1927 |
| p | 0.0154 | 0.2875 | 0.6442 |
| | * | ns | ns |
| | Exact | Exact | Exact |

**Supplemental Figure 9: Correlation between MOS-specific IgG and disease outcomes.** Scatterplot depicting the magnitude of MOS-specific IgG responses at the post-immunization timepoint for animals from Darrah et al., Nature 2020. Route and dose group are indicated in symbol shape and color. Spearman correlation coefficient ($R_S$) and p value for MOS IgG versus FDG and CFU disease outcomes are reported in inset across all animals in the study, and tabulated below within individual dose and route groups.

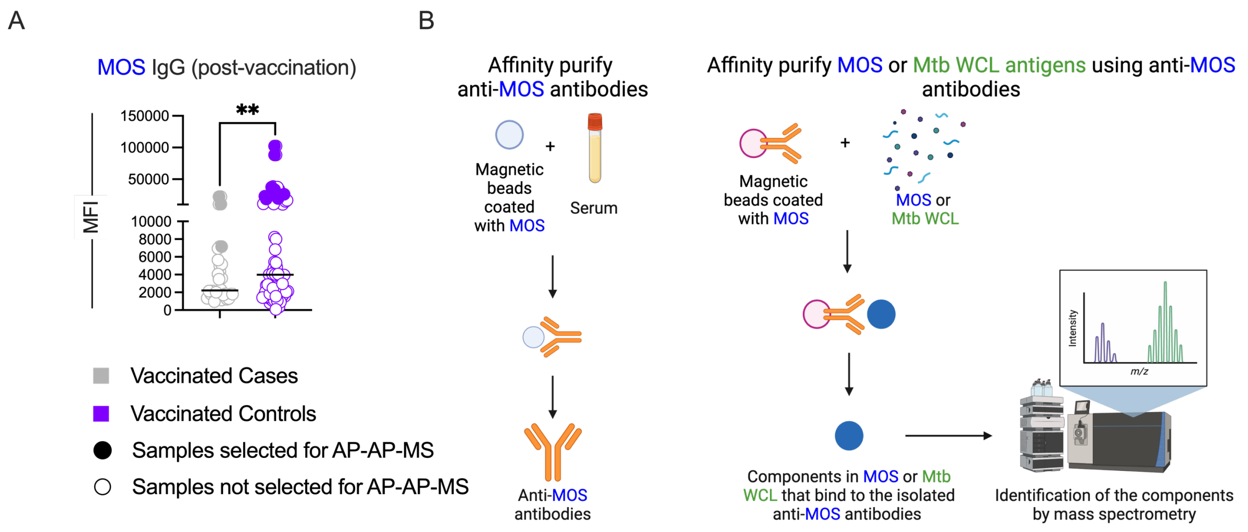

**Supplemental Figure 10: Schematic for immunoproteomics. A.** Box plots of MOS IgG in vaccinated controls and cases at post-vaccination timepoint demonstrating select serum samples that were used to isolate MOS-specific antibodies (solid). **B.** Isolating immunogenic peptides from MOS or Mtb WCL that bind to anti-MOS antibodies was a two-step process. The first step (left) included enriching anti-MOS antibodies from serum. For that, magnetic beads coated with MOS were incubated with serum, and the components that bind to the beads (anti-MOS antibodies) were eluted. The eluted anti-MOS antibodies were then conjugated on magnetic beads (right), which were incubated with MOS or Mtb WCL. Components in the MOS or Mtb WCL that bind to the MOS-specific antibodies were then eluted and detected by mass spectrometry.

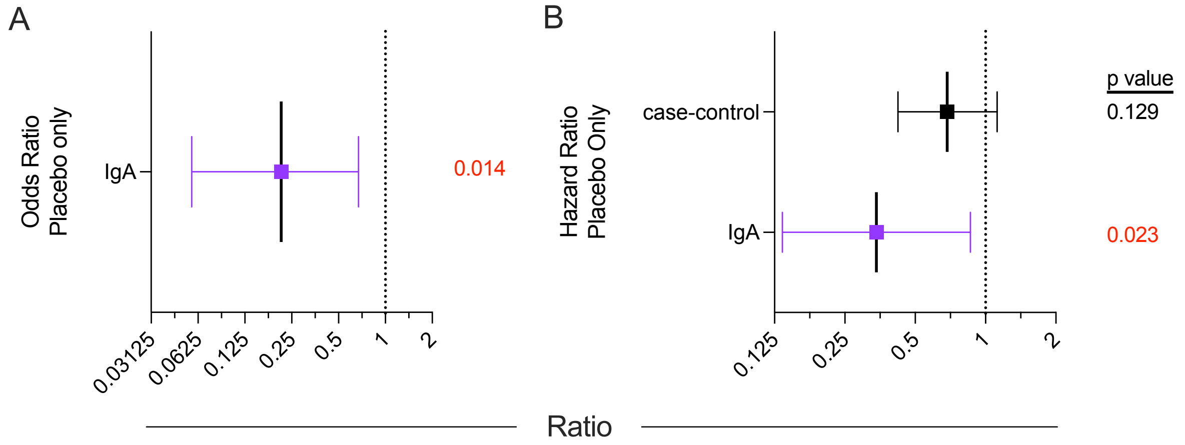

**Supplemental Figure 11: Placebo group correlate analysis**. **A-B**. Odds ratio (**A**) and Hazard Ratio (**B**) of TB disease per 10-fold increase in each MOS-specific IgA. Statistical significance (p value) is indicated in inset. Values < 0.1 are shown in red. The case-control hazard ratio indicates that observed among the subset of participants selected for case-control dataset, independently of immune markers.

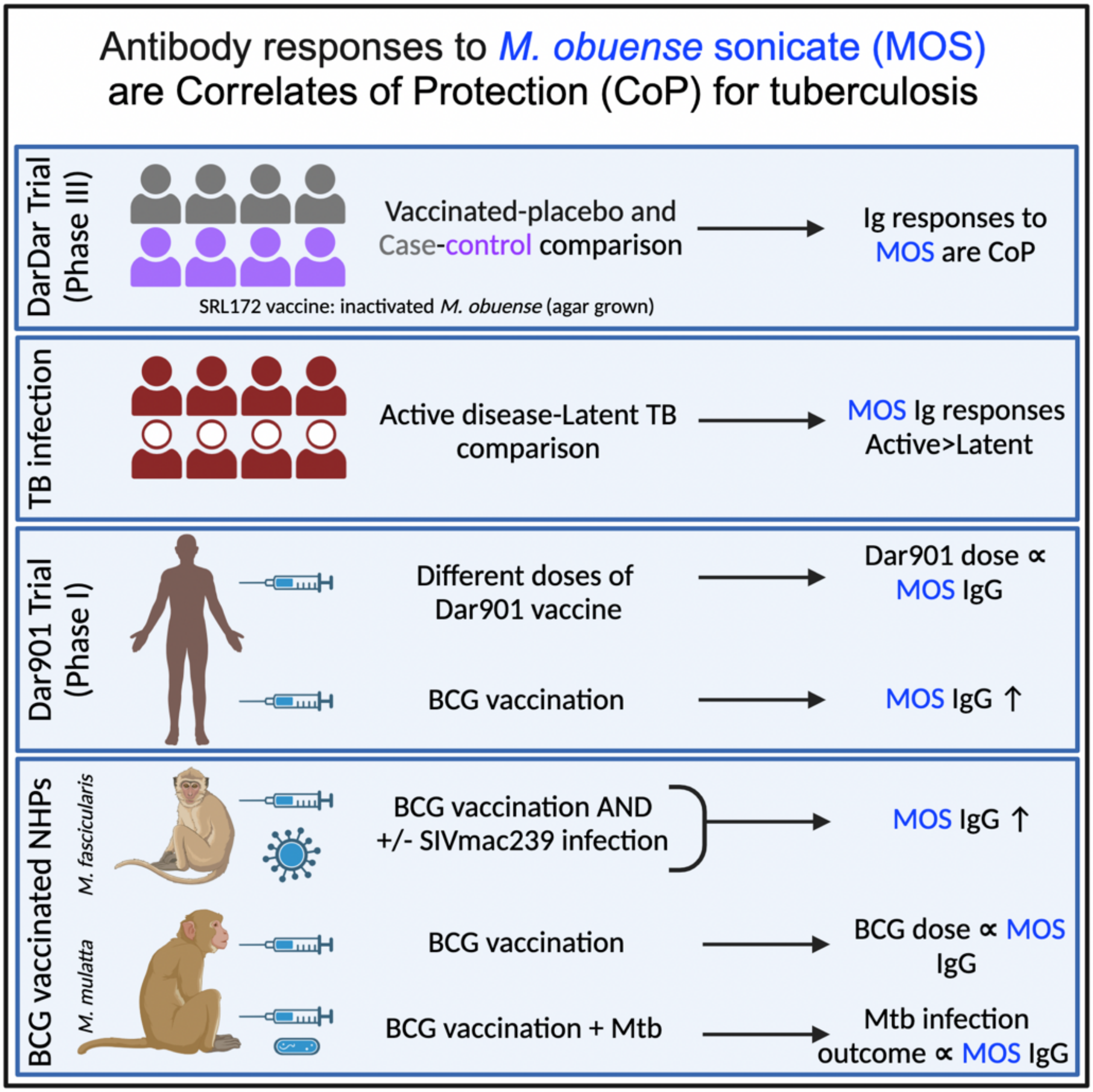

**Supplemental Figure 12: Summary of cohorts used and key observations.**

**Supplemental Table 1: Characteristics of case control cohort of subjects from DarDar trial:** The table highlights the meta-data from participants of DarDar trial, a phase III clinical trial conducted in Tanzania in people living with HIV-1, that had CD4 counts of at least 200 cells/μL and a BCG scar, and received 5 doses of 1 mg inactivated *M. obuense* SRL172 or borate-buffered isotonic saline intradermally.

| | Vaccinated (n=95) | | Placebo (n=105) | |
|---|---|---|---|---|
| | Control | Case | Control | Case |
| **Number (n)** | 68 | 27 | 64 | 41 |
| **Average age (s.d.)** | 34 (7.85) | 38 (8.40) | 35 (8.24) | 35 (7.40) |
| **Sex** | | | | |
| Male | 16 | 8 | 15 | 8 |
| Female | 52 | 19 | 49 | 33 |
| **Average Pre-vaccination CD4+ Count** | 397 (168.40) | 399 (164.93) | 390 (163.57) | 407 (179.00) |
| **PPD test result** | | | | |
| Positive | 40 | 13 | 38 | 21 |
| Negative | 28 | 14 | 26 | 20 |
| **Previous TB infection** | | | | |
| Yes | 10 | 4 | 10 | 8 |
| No | 58 | 23 | 54 | 33 |

**Supplemental Table 2: Antigens used for Fc Array Assay**

| ***Mycobacterium obuense* antigens** | | |
|---|---|---|
| Antigen | | Source |
| MOS [*M. obuense* sonicate (agar grown)] | | Dartmouth Health |
| MO WCL [*M. obuense* WCL (broth grown)] | | Dartmouth Health |
| MOM [inactivated *M. obuense* mycobacteria (SRL172 vaccine)] | | Dartmouth Health |
| ***Mycobacterium tuberculosis* antigens** | | |
| Antigen | Antigen production details | Source |
| Mtb Ag85 | Purified from Mtb H37Rv | BEI, NR-14855 |
| Mtb APA (Mpt32) | Purified from Mtb H37Rv | BEI, NR-14862 |
| Mtb Crystallin | Purified from Mtb H37Rv | BEI, NR-14860 |
| Mtb Esat6 | Recombinant protein (*E. coli* expression) | BEI, NR-14868 |
| Mtb GroES | Purified from Mtb H37Rv | BEI, NR-14861 |
| Mtb LAM | Purified from Mtb H37Rv | BEI, NR-53528 |
| Mtb Mpt64 | Recombinant protein (*E. coli* expression) | BEI, NR-49435 |
| Mtb PstS1 | Purified from Mtb H37Rv | BEI, NR-53528 |
| Mtb CDC1551 WCL | WCL from Mtb CDC1551 | BEI, NR-14823 |
| Mtb 91_0079 WCL | WCL from Mtb East African Indian 91_0079 | BEI, NR-36497 |
| Mtb H37Rv Cytosol | Cytosol fraction from Mtb H37Rv | BEI, NR-14834 |
| Mtb H37Rv Membrane | Cell membrane fraction from Mtb H37Rv | BEI, NR-14831 |
| Mtb H37Rv Filtrate | Mtb H37Rv culture filtrate | BEI, NR-14825 |

**Supplemental Table 3: Identification of vaccine-induced antibody responses.** Antibody response features induced by SRL172 immunization (placebo vs SRL172 recipient), as defined as those that exhibited p<0.10 when analyzed in the context of multiple hypothesis correction (False Discovery Rate with Benjamini Hochberg method) at pre- and post-immunization timepoints. P values less than 0.1 following MHC are indicated in red. Features are ordered by post-immunization timepoint p value.

| Feature | Pre-immunization | | Post-immunization | |
|---|---|---|---|---|
| | p-value | Adjusted p-value | p-value | Adjusted p-value |
| MOS Fcar | 0.44260181 | 0.87249186 | 5.32E-07 | 9.20E-05 |
| MOS Fcgr2a | 0.68814259 | 0.98349362 | 2.35E-06 | 0.00020338 |
| MOS Fcgr3a | 0.47599664 | 0.89095807 | 5.36E-06 | 0.00030918 |
| MOS IgG | 0.22182033 | 0.8543905 | 6.33E-05 | 0.0027388 |
| MOS IgA | 0.28928342 | 0.8543905 | 0.00011237 | 0.00333864 |
| MOS Fcgr2b | 0.43336091 | 0.87249186 | 0.00011579 | 0.00333864 |
| MOS IgG2 | 0.38828638 | 0.8543905 | 0.00019647 | 0.00485557 |
| MOS Fcgr3b | 0.3537092 | 0.8543905 | 0.00099697 | 0.02155957 |
| MOS IgG1 | 0.29809007 | 0.8543905 | 0.00391109 | 0.07517977 |
| Mtb Esat6 IgG4 | 0.98761126 | 0.99738739 | 0.01051143 | 0.18184765 |
| Mtb Mpt64 IgG | 0.35494826 | 0.8543905 | 0.02694391 | 0.42375422 |
| Mtb H37Rv Culture Filtrate Fraction _IgM | 0.00814063 | 0.54194863 | 0.03226162 | 0.44093205 |
| Mtb GroES IgM | 0.35166805 | 0.8543905 | 0.03313362 | 0.44093205 |
| Mtb LAM Fcgr2b | 0.95016604 | 0.99738739 | 0.04498254 | 0.53791338 |
| Mtb CDC1551 IgM | 0.0561511 | 0.6476093 | 0.05076521 | 0.53791338 |
| Mtb LAM IgG | 0.9047586 | 0.99738739 | 0.05144374 | 0.53791338 |
| MOS IgG3 | 0.87436158 | 0.99738739 | 0.05285854 | 0.53791338 |
| Mtb Mpt64 IgG1 | 0.42313645 | 0.87249186 | 0.05854275 | 0.56266089 |
| Mtb 91_0079 WCL IgG | 0.78665321 | 0.98349362 | 0.07286646 | 0.6634683 |
| Mtb Mpt64 IgG2 | 0.34207012 | 0.8543905 | 0.076953 | 0.66564349 |
| Mtb Esat6 IgA | 0.29796571 | 0.8543905 | 0.08495017 | 0.69982759 |
| Mtb CDC1551 Fcgr2a | 0.29209629 | 0.8543905 | 0.10100475 | 0.76204793 |
| Mtb H37Rv Cytosol Fraction FcgR3b | 0.99608333 | 0.99738739 | 0.11179882 | 0.76204793 |
| Mtb H37Rv Cell Membrane Fraction IgG1 | 0.07706975 | 0.70174034 | 0.11334619 | 0.76204793 |
| Mtb CDC1551 Fcgr3b | 0.63355728 | 0.97688764 | 0.11490353 | 0.76204793 |
| Mtb CDC1551 Fcgr2b | 0.99738739 | 0.99738739 | 0.11712013 | 0.76204793 |
| Mtb H37Rv Cell Membrane Fraction IgG3 | 0.68981164 | 0.98349362 | 0.11943836 | 0.76204793 |
| Mtb 91_0079 WCL IgM | 0.05040267 | 0.6476093 | 0.12333724 | 0.76204793 |
| Mtb 91_0079 WCL FcgR3b | 0.21650183 | 0.8543905 | 0.13030923 | 0.76993937 |
| Mtb H37Rv Cytosol Fraction IgM | 0.03432531 | 0.54194863 | 0.13613955 | 0.76993937 |
| Mtb Mpt64 IgG3 | 0.29625214 | 0.8543905 | 0.14863156 | 0.76993937 |
| Mtb Mpt64 FcgR2a | 0.89719557 | 0.99738739 | 0.14916311 | 0.76993937 |
| Mtb CDC1551 IgG3 | 0.31019528 | 0.8543905 | 0.15087506 | 0.76993937 |
| Mtb Ag85 Fcar | 0.76192299 | 0.98349362 | 0.15432392 | 0.76993937 |
| Mtb LAM IgG3 | 0.75079005 | 0.98349362 | 0.15711102 | 0.76993937 |
| Mtb Crystallin IgA | 0.51261018 | 0.90670379 | 0.1602186 | 0.76993937 |
| Mtb 91_0079 WCL FcgR3a | 0.86750596 | 0.99738739 | 0.1828496 | 0.77221759 |
| Mtb CDC1551 IgA | 0.4538975 | 0.87249186 | 0.18329934 | 0.77221759 |
| Mtb 91_0079 WCL FcgR2a | 0.795869 | 0.98349362 | 0.18679039 | 0.77221759 |
| Mtb Esat6 IgM | 0.00672972 | 0.54194863 | 0.198184 | 0.77221759 |
| Mtb Ag85 IgA | 0.80157573 | 0.98349362 | 0.20276583 | 0.77221759 |
| MO WCL IgA | 0.45230039 | 0.87249186 | 0.20376521 | 0.77221759 |
| MO WCL IgG2 | 0.0247009 | 0.54194863 | 0.20921002 | 0.77221759 |
| Mtb Mpt64 FcgR2b | 0.9593343 | 0.99738739 | 0.21512382 | 0.77221759 |
| MOS IgM | 0.02831885 | 0.54194863 | 0.22525881 | 0.77221759 |
| Mtb Esat6 IgG | 0.65313417 | 0.98254097 | 0.22791558 | 0.77221759 |
| Mtb LAM Fcgr2a | 0.83021018 | 0.99738739 | 0.22819879 | 0.77221759 |
| Mtb GroES IgG4 | 0.22195201 | 0.8543905 | 0.22843331 | 0.77221759 |
| Mtb APA IgM | 0.11855293 | 0.82038628 | 0.23495537 | 0.77221759 |
| Mtb Mpt64 FcgR3a | 0.65164323 | 0.98254097 | 0.23798114 | 0.77221759 |
| Mtb LAM IgG2 | 0.89135805 | 0.99738739 | 0.23819224 | 0.77221759 |
| Mtb Mpt64 IgG4 | 0.19290175 | 0.8543905 | 0.24150841 | 0.77221759 |
| Mtb CDC1551 Fcar | 0.29421399 | 0.8543905 | 0.25022204 | 0.77221759 |
| Mtb Esat6 IgG2 | 0.02047287 | 0.54194863 | 0.25412279 | 0.77221759 |
| Mtb Esat6 IgG1 | 0.19236841 | 0.8543905 | 0.25701972 | 0.77221759 |

| | | | | |
|---|---|---|---|---|
| Mtb Crystallin IgG2 | 0.26498056 | 0.8543905 | 0.26509881 | 0.77221759 |
| Mtb APA IgA | 0.27769484 | 0.8543905 | 0.27275404 | 0.77221759 |
| Mtb APA IgG2 | 0.04506583 | 0.6476093 | 0.27528618 | 0.77221759 |
| Mtb H37Rv Cytosol Fraction IgG3 | 0.26573455 | 0.8543905 | 0.28163113 | 0.77221759 |
| Mtb PstS1 IgM | 0.08414145 | 0.72415757 | 0.29231269 | 0.77221759 |
| Mtb CDC1551 IgG2 | 0.51362411 | 0.90670379 | 0.2934723 | 0.77221759 |
| MO WCL Fcar | 0.39758625 | 0.8543905 | 0.2941415 | 0.77221759 |
| Mtb H37Rv Cell Membrane Fraction IgG2 | 0.55284231 | 0.9224727 | 0.2947071 | 0.77221759 |
| Mtb H37Rv Cell Membrane Fraction FcgR3b | 0.73443326 | 0.98349362 | 0.29776269 | 0.77221759 |
| Mtb H37Rv Culture Filtrate Fraction IgA | 0.47895434 | 0.89095807 | 0.30074118 | 0.77221759 |
| MO WCL IgM | 0.01524845 | 0.54194863 | 0.30081272 | 0.77221759 |
| Mtb GroES IgG1 | 0.91825647 | 0.99738739 | 0.30094937 | 0.77221759 |
| Mtb 91_0079 WCL IgG2 | 0.94790907 | 0.99738739 | 0.31412517 | 0.77221759 |
| Mtb CDC1551 Fcgr3a | 0.19783183 | 0.8543905 | 0.32019768 | 0.77221759 |
| Mtb Crystallin Fcgr3b | 0.43862842 | 0.87249186 | 0.32190138 | 0.77221759 |
| Mtb Ag85 IgG4 | 0.37918288 | 0.8543905 | 0.32294062 | 0.77221759 |
| Mtb 91_0079 WCL IgG4 | 0.75429922 | 0.98349362 | 0.32410204 | 0.77221759 |
| Mtb H37Rv Culture Filtrate Fraction .Protein_IgG | 0.5816732 | 0.94046228 | 0.32584904 | 0.77221759 |
| Mtb H37Rv Cytosol Fraction Fcar | 0.36003433 | 0.8543905 | 0.33248302 | 0.77634538 |
| Mtb PstS1 IgG | 0.725649 | 0.98349362 | 0.33864985 | 0.77634538 |
| Mtb APA IgG1 | 0.33772371 | 0.8543905 | 0.34105346 | 0.77634538 |
| Mtb PstS1 IgG1 | 0.74120493 | 0.98349362 | 0.3460738 | 0.77754244 |
| Mtb H37Rv Culture Filtrate Fraction .Protein_IgG3 | 0.1923337 | 0.8543905 | 0.35360593 | 0.78427981 |
| Mtb LAM IgM | 0.78856734 | 0.98349362 | 0.37218567 | 0.80698122 |
| Mtb GroES IgG3 | 0.84486345 | 0.99738739 | 0.37317051 | 0.80698122 |
| Mtb 91_0079 WCL Fcar | 0.30125106 | 0.8543905 | 0.38377142 | 0.80727423 |
| Mtb 91_0079 WCL FcgR2b | 0.63808268 | 0.97688764 | 0.39116353 | 0.80727423 |
| Mtb H37Rv Cytosol Fraction IgA | 0.40003255 | 0.8543905 | 0.39470525 | 0.80727423 |
| Mtb PstS1 IgG2 | 0.03445916 | 0.54194863 | 0.39632787 | 0.80727423 |
| Mtb LAM Fcgr3a | 0.95137546 | 0.99738739 | 0.40009332 | 0.80727423 |
| Mtb GroES FcgR3a | 0.72705265 | 0.98349362 | 0.40337797 | 0.80727423 |
| Mtb H37Rv Culture Filtrate Fraction .Protein_IgG1 | 0.5366731 | 0.9224727 | 0.40597028 | 0.80727423 |
| Mtb Ag85 IgG1 | 0.29912796 | 0.8543905 | 0.42011727 | 0.81034191 |
| Mtb LAM IgG4 | 0.55610365 | 0.9224727 | 0.42634696 | 0.81034191 |
| Mtb Mpt64 FcgR3b | 0.93070707 | 0.99738739 | 0.42845581 | 0.81034191 |
| Mtb CDC1551 IgG4 | 0.05611047 | 0.6476093 | 0.43131845 | 0.81034191 |
| Mtb H37Rv Cytosol Fraction IgG2 | 0.93180628 | 0.99738739 | 0.43199008 | 0.81034191 |
| Mtb Crystallin IgG | 0.0882295 | 0.72415757 | 0.4360282 | 0.81034191 |
| Mtb PstS1 Fcgr2b | 0.31929678 | 0.8543905 | 0.44030138 | 0.81034191 |
| Mtb Ag85 IgG3 | 0.34965999 | 0.8543905 | 0.44577531 | 0.8117803 |
| Mtb GroES IgG2 | 0.92105998 | 0.99738739 | 0.45403317 | 0.81820561 |
| Mtb H37Rv Culture Filtrate Fraction _IgG4 | 0.11120715 | 0.80161818 | 0.47890238 | 0.84260758 |
| Mtb GroES FcgR2b | 0.63546168 | 0.97688764 | 0.48042933 | 0.84260758 |
| Mtb H37Rv Cell Membrane Fraction FcgR2b | 0.06788023 | 0.70174034 | 0.48218584 | 0.84260758 |
| Mtb GroES FcgR3b | 0.96699366 | 0.99738739 | 0.49182386 | 0.84818354 |
| Mtb PstS1 Fcgr3b | 0.339812 | 0.8543905 | 0.49739624 | 0.84818354 |
| MO WCL Fcgr2a | 0.09627528 | 0.72415757 | 0.50117863 | 0.84818354 |
| Mtb Crystallin IgG4 | 0.16770123 | 0.8543905 | 0.5066422 | 0.84818354 |
| Mtb LAM Fcgr3b | 0.72084091 | 0.98349362 | 0.50989068 | 0.84818354 |
| Mtb H37Rv Cell Membrane Fraction IgG | 0.99197234 | 0.99738739 | 0.51863266 | 0.85450905 |
| Mtb H37Rv Cytosol Fraction FcgR3a | 0.39727786 | 0.8543905 | 0.53210104 | 0.86515058 |
| Mtb H37Rv Culture Filtrate Fraction IgG2 | 0.02889162 | 0.54194863 | 0.53793094 | 0.86515058 |
| Mtb Ag85 IgG | 0.32682681 | 0.8543905 | 0.54040798 | 0.86515058 |
| Mtb H37Rv Cytosol Fraction IgG4 | 0.51235863 | 0.90670379 | 0.54509487 | 0.86515058 |
| Mtb 91_0079 WCL IgG1 | 0.39909537 | 0.8543905 | 0.57758029 | 0.90222127 |
| Mtb GroES IgG | 0.94672926 | 0.99738739 | 0.57888185 | 0.90222127 |
| Mtb APA IgG3 | 0.36923263 | 0.8543905 | 0.59122919 | 0.91323794 |
| Mtb 91_0079 WCL IgG3 | 0.32017968 | 0.8543905 | 0.60392606 | 0.92459476 |
| Mtb H37Rv Culture Filtrate Fraction .Protein_Fcar | 0.24540591 | 0.8543905 | 0.6150296 | 0.93083102 |
| Mtb Crystallin Fcgr3a | 0.94982923 | 0.99738739 | 0.6187605 | 0.93083102 |
| Mtb GroES Fcar | 0.09468914 | 0.72415757 | 0.62845304 | 0.93395591 |
| MOS IgG4 | 0.43338506 | 0.87249186 | 0.64605475 | 0.93395591 |
| Mtb H37Rv Cytosol Fraction FcgR2b | 0.55499708 | 0.9224727 | 0.65385046 | 0.93395591 |

| | | | | |
|---|---|---|---|---|
| Mtb CrystallinFcar | 0.30156398 | 0.8543905 | 0.66818259 | 0.93395591 |
| Mtb H37Rv Culture Filtrate Fraction .Protein_FcgR2b | 0.53854316 | 0.9224727 | 0.66821934 | 0.93395591 |
| Mtb H37Rv Culture Filtrate Fraction FcgR3a | 0.24044468 | 0.8543905 | 0.67152453 | 0.93395591 |
| Mtb Ag85 IgG2 | 0.37844131 | 0.8543905 | 0.67269757 | 0.93395591 |
| Mtb H37Rv Cell Membrane Fraction IgG4 | 0.14513622 | 0.8543905 | 0.67329885 | 0.93395591 |
| Mtb CDC1551 IgG | 0.88097927 | 0.99738739 | 0.67642069 | 0.93395591 |
| Mtb LAM Fcar | 0.93921642 | 0.99738739 | 0.68606672 | 0.93395591 |
| Mtb Mpt64 IgM | 0.55988227 | 0.9224727 | 0.68640219 | 0.93395591 |
| Mtb APA Fcgr2b | 0.70390408 | 0.98349362 | 0.68891059 | 0.93395591 |
| Mtb CDC1551 IgG1 | 0.26426242 | 0.8543905 | 0.69668152 | 0.93395591 |
| Mtb Mpt64 Fcar | 0.78471271 | 0.98349362 | 0.70586488 | 0.93395591 |
| Mtb APA Fcgr3a | 0.85797089 | 0.99738739 | 0.7120998 | 0.93395591 |
| Mtb Ag85 FcgR3a | 0.39975221 | 0.8543905 | 0.71274497 | 0.93395591 |
| Mtb APA Fcar | 0.17016394 | 0.8543905 | 0.71591376 | 0.93395591 |
| Mtb H37Rv Cytosol Fraction IgG1 | 0.38455602 | 0.8543905 | 0.71919728 | 0.93395591 |
| Mtb H37Rv Cell Membrane Fraction IgA | 0.99303468 | 0.99738739 | 0.72945016 | 0.93395591 |
| Mtb LAM IgA | 0.49179194 | 0.895579 | 0.73363094 | 0.93395591 |
| Mtb H37Rv Cell Membrane Fraction Fcar | 0.89636289 | 0.99738739 | 0.73576035 | 0.93395591 |
| Mtb Mpt64 IgA | 0.35997632 | 0.8543905 | 0.7396067 | 0.93395591 |
| Mtb Crystallin IgG3 | 0.44454934 | 0.87249186 | 0.75306294 | 0.94405717 |
| MO WCL IgG | 0.67320538 | 0.98349362 | 0.76289901 | 0.9495074 |
| Mtb H37Rv Culture Filtrate Fraction .Protein_FcgR3b | 0.79104944 | 0.98349362 | 0.78491641 | 0.96662077 |
| Mtb 91_0079 WCL IgA | 0.93314105 | 0.99738739 | 0.78782387 | 0.96662077 |
| Mtb PstS1 IgA | 0.61268247 | 0.97688764 | 0.80278759 | 0.96700051 |
| MO WCL IgG1 | 0.95101608 | 0.99738739 | 0.81120226 | 0.96700051 |
| Mtb H37Rv Cell Membrane Fraction FcgR3a | 0.39041904 | 0.8543905 | 0.82567488 | 0.96700051 |
| Mtb APA Fcgr2a | 0.8256604 | 0.99738739 | 0.82970748 | 0.96700051 |
| MO WCL IgG4 | 0.70593683 | 0.98349362 | 0.8299006 | 0.96700051 |
| Mtb Esat6 Fcgr3a | 0.98854511 | 0.99738739 | 0.83168247 | 0.96700051 |
| Mtb Ag85 FcgR3b | 0.46631034 | 0.88650208 | 0.83275546 | 0.96700051 |
| MO WCL Fcgr3b | 0.43095565 | 0.87249186 | 0.83285015 | 0.96700051 |
| Mtb Crystallin Fcgr2a | 0.61890575 | 0.97688764 | 0.83934478 | 0.96804432 |
| Mtb PstS1 Fcgr3a | 0.7882982 | 0.98349362 | 0.84918819 | 0.97291097 |
| Mtb Esat6 Fcar | 0.28059874 | 0.8543905 | 0.85964973 | 0.97841712 |
| Mtb APA IgG | 0.32704338 | 0.8543905 | 0.86715948 | 0.98051366 |
| Mtb PstS1 IgG4 | 0.01344032 | 0.54194863 | 0.88653701 | 0.99239399 |
| Mtb PstS1 Fcar | 0.07638636 | 0.70174034 | 0.8949643 | 0.99239399 |
| Mtb Esat6 Fcgr2a | 0.88005004 | 0.99738739 | 0.89714708 | 0.99239399 |
| Mtb GroES IgA | 0.27641837 | 0.8543905 | 0.90121582 | 0.99239399 |
| Mtb PstS1 IgG3 | 0.54131909 | 0.9224727 | 0.91401046 | 0.99239399 |
| Mtb H37Rv Cytosol Fraction IgG | 0.56731356 | 0.92589854 | 0.92023443 | 0.99239399 |
| Mtb PstS1 Fcgr2a | 0.07695205 | 0.70174034 | 0.92485069 | 0.99239399 |
| Mtb Esat6 IgG3 | 0.29334474 | 0.8543905 | 0.92828515 | 0.99239399 |
| MO WCL IgG3 | 0.73715576 | 0.98349362 | 0.93572858 | 0.99239399 |
| Mtb Esat6 Fcgr3b | 0.48951844 | 0.895579 | 0.94488222 | 0.99239399 |
| Mtb Crystallin IgG1 | 0.79867743 | 0.98349362 | 0.94925094 | 0.99239399 |
| Mtb H37Rv Cell Membrane Fraction IgM | 0.78493845 | 0.98349362 | 0.95608303 | 0.99239399 |
| Mtb Ag85 IgM | 0.02642003 | 0.54194863 | 0.96621405 | 0.99239399 |
| Mtb LAM IgG1 | 0.67419035 | 0.98349362 | 0.9701376 | 0.99239399 |
| Mtb Esat6 Fcgr2b | 0.6348457 | 0.97688764 | 0.97118061 | 0.99239399 |
| Mtb APA Fcgr3b | 0.67592791 | 0.98349362 | 0.97723231 | 0.99239399 |
| MO WCL Fcgr3a | 0.29012489 | 0.8543905 | 0.97972505 | 0.99239399 |
| Mtb APA IgG4 | 0.75363666 | 0.98349362 | 0.98256549 | 0.99239399 |
| Mtb Ag85 FcgR2b | 0.31241576 | 0.8543905 | 0.98665761 | 0.99239399 |
| MO WCL Fcgr2b | 0.38102466 | 0.8543905 | 0.99920869 | 0.99920869 |

**Supplemental Table 4: Case-control analysis of vaccine-induced immune responses.** Features induced by immunization in the vaccinated group, defined as those that exhibited p<0.1, were analyzed in the context of multiple hypothesis correction (False Discovery Rate with Benjamini Hochberg method) in the vaccinated group at post-vaccination timepoint. P values less than 0.1 after MHC are indicated in red.

| Feature induced by immunization | Unadjusted p-value | Adjusted p-value |
|---|---|---|
| MOS IgG | 0.00951 | 0.0374 |
| MOS FcgR2A | 0.0105 | 0.0374 |
| MOS FcgR3A | 0.0145 | 0.0374 |
| MOS IgA | 0.0166 | 0.0374 |
| MOS FcgR2B | 0.0361 | 0.0547 |
| MOS IgG1 | 0.0365 | 0.0547 |
| MOS FcgR3B | 0.0794 | 0.102 |
| MOS IgG2 | 0.135 | 0.152 |
| MOS FcaR | 0.257 | 0.257 |

**Supplemental Table 5: DAR-901 trial study cohort:** The table highlights meta-data from DAR-901 Phase I dose escalation trial conducted in US-based participants.

| | DAR-901 | | | BCG | Placebo |
|---|---|---|---|---|---|
| | 0.1 mg | 0.3 mg | 1.0 mg | | |
| **Number (n)** | 10 | 10 | 20 | 9 | 9 |
| **Average Age (s.d.)** | 41.00 (14.88) | 27.90 (12.41) | 36.6 (13.80) | 30.00 (11.81) | 30.00 (8.97) |
| **Sex** | | | | | |
| Male | 5 | 3 | 9 | 4 | 4 |
| Female | 5 | 7 | 11 | 5 | 5 |
| **Ethnicity** | | | | | |
| Hispanic or Latino | 0 | 2 | 3 | 0 | 0 |
| Non- Hispanic or Latino | 10 | 8 | 17 | 9 | 9 |
| **Race** | | | | | |
| Black or African American | 0 | 1 | 4 | 4 | 1 |
| White | 5 | 7 | 8 | 3 | 5 |
| Asian | 5 | 2 | 8 | 2 | 3 |

**Supplemental Table 6: TB disease and TB infection samples cohort:** The table highlights meta-data in people diagnosed with TB disease and TB infection acquired from Duke University.

| | TB disease (Active TB) | | TB infection (Latent TB) |
|---|---|---|---|
| | Clinical | Culture proven | |
| **Number (n)** | 3 | 22 | 25 |
| **Sex** | | | |
| Male | 3 | 18 | 18 |
| Female | 0 | 4 | 7 |
| **Place of birth** | | | |
| Australia | 0 | 0 | 1 |
| El Salvador | 0 | 1 | 2 |
| Ethiopia | 0 | 0 | 1 |
| Gambia | 0 | 0 | 1 |
| Honduras | 0 | 1 | 1 |
| India | 0 | 3 | 2 |
| Liberia | 0 | 1 | 0 |
| Mexico | 1 | 2 | 5 |
| Morocco | 0 | 2 | 0 |
| Nigeria | 0 | 1 | 1 |
| USA | 2 | 10 | 11 |
| Zimbabwe | 0 | 1 | 0 |
| **Race** | | | |
| Asian | 0 | 3 | 2 |
| Black | 0 | 10 | 11 |
| White | 3 | 9 | 12 |
| **Ethnicity** | | | |
| Hispanic | 1 | 6 | 8 |
| Non-Hispanic | 2 | 15 | 17 |
| Unknown | 0 | 1 | 0 |
| **HIV status** | | | |
| Positive | 2 | 19 | 21 |
| Negative | 1 | 3 | 2 |
| Unknown | 0 | 0 | 2 |
| Previous TB infection | | | |
| Yes | 2 | 9 | 0 |
| No | 1 | 12 | 24 |
| Unknown | 0 | 1 | 1 |

**Supplemental Table 7: Non-human Primate cohorts.**

| Study | Monkey Species | Study description | Groups | Sample size |
|---|---|---|---|---|
| Jauro et al, 2024 | *Macaca fascicularis* | Animals infected with SIV and administered BCG using different routes | i.v. BCG (SIV infected) | 5 |
| | | | i.d. BCG (SIV infected) | 6 |
| Larson et al, 2023 | *Macaca fascicularis* | Animals administered with i.v. BCG with or without prior SIV infection, and challenged with Mtb | i.v. BCG (SIV naïve) | 7 |
| | | | i.v. BCG (SIV infected) | 9 |
| Darrah et al, 2023 | *Macaca mulatta* | SIV naïve animals administered with varying dose of BCG, and infected with Mtb | 4.5-6.0 $\log_{10}$CFU i.v. BCG | 18 |
| | | | 6.0-7.5 $\log_{10}$CFU i.v. BCG | 16 |
| | | | Unvaccinated | 1 |
| Darrah et al, 2020 | *Macaca mulatta* | SIV naïve animals vaccinated with BCG (varying route and dose), and infected with Mtb | i.d. low ($5 \times 10^5$ CFU) | 10 |
| | | | i.d. high ($5 \times 10^7$ CFU) | 8 |
| | | | i.v. BCG ($5 \times 10^7$ CFU) | 10 |
| | | | Unvaccinated | 4 |

### Supplemental Table 8: MOS proteins pulled down by pre-immunization serum antibodies.

| MOS protein [Uniprot ID] | Top 3 BLAST hits on Mtb proteome | | | Proteomic Identification |
|---|---|---|---|---|
| | Mtb strain | Protein name [Uniprot ID] | % Similarity | |
| Ketol-acid reductoisomerase (NADP(+)) [A0A0M2K621] | ATCC25618/ H37Rv | Ketol-acid reductoisomerase (NADP(+)) [P9WKJ7] | 95 | Ketol-acid reductoisomerase (NADP(+)) OS=Mycolicibacterium obuense OX=1807 GN=ilvC PE=3 SV=1; Ketol-acid reductoisomerase (NADP(+)) OS=Mycolicibacterium obuense OX=1807 GN=ilvC PE=3 SV=1 |
| | CDC1551/ Oshkosh | Ketol-acid reductoisomerase (NADP(+)) [P9WKJ6] | 95 | |
| | ATCC25177/ H37Ra | Ketol-acid reductoisomerase (NADP(+)) [A5U713] | 95 | Ketol-acid reductoisomerase (NADP(+)) OS=Mycolicibacterium obuense OX=1807 GN=ilvC PE=3 SV=1 |
| PspA/IM30 family protein [A0A0J6W046] | ATCC25618/ H37Rv | PspA protein [P9WHP5] | 89.5 | PspA/IM30 family protein OS=Mycolicibacterium obuense OX=1807 GN=EUA04_24980 PE=3 SV=1 |
| | CDC1551/ Oshkosh | PspA protein [P9WHP4] | 89.5 | |
| | ATCC25177/ H37Ra | PspA protein [A5U697] | 89.5 | |
| ABC transporter substrate-binding protein [A0A0M2JQE2] | ATCC25618/ H37Rv | Probable glutamine-binding lipoprotein GlnH (GLNBP) [P96257] | 37.2 | ABC transporter substrate-binding protein OS=Mycolicibacterium obuense OX=1807 GN=WN67_27560 PE=4 SV=1; ABC transporter glutamine-binding protein GlnH OS=Mycolicibacterium obuense OX=1807 GN=glnH_4 PE=4 SV=1 |
| | CDC1551/ Oshkosh | Amino acid ABC transporter, amino acid-binding protein [L7N636] | 37.2 | |
| | ATCC25177/ H37Ra | Amino acid ABC transporter substrate-binding protein [A5TZD7] | 37.2 | |
| Probable cytosol aminopeptidase [A0A0J6YPN8] | ATCC25618/ H37Rv | Probable cytosol aminopeptidase [P9WHT3] | 81.8 | Probable cytosol aminopeptidase OS=Mycolicibacterium obuense OX=1807 GN=pepA_2 PE=3 SV=1 |
| | CDC1551/ Oshkosh | Probable cytosol aminopeptidase [P9WHT2] | 81.8 | |
| | ATCC25177/ H37Ra | Probable cytosol aminopeptidase [A5U4P1] | 81.8 | |
| Glyceraldehyde-3-phosphate dehydrogenase [A0A0J6WC82] | ATCC25618/ H37Rv | Glyceraldehyde-3-phosphate dehydrogenase [P9WN83] | 92.9 | glyceraldehyde-3-phosphate dehydrogenase [Source:HGNC Symbol;Acc:4141]; Glyceraldehyde-3-phosphate dehydrogenase OS=Mycolicibacterium obuense OX=1807 GN=gapA PE=3 SV=1; SWISS-PROT:P10096 (Bos taurus) Glyceraldehyde-3-phosphate dehydrogenase |
| | CDC1551/ Oshkosh | Glyceraldehyde-3-phosphate dehydrogenase [P9WN82] | 92.9 | |
| | ATCC25177/ H37Ra | Glyceraldehyde-3-phosphate dehydrogenase [A5U2D8] | 92.9 | |
| Superoxide dismutase [A0A0M2JQV1] | ATCC25618/ H37Rv | Superoxide dismutase (Fe) [P9WGE7] | 86 | Superoxide dismutase OS=Mycolicibacterium obuense OX=1807 GN=sodA PE=3 SV=1; Superoxide dismutase OS=Mycolicibacterium obuense OX=1807 GN=WN67_26715 PE=3 SV=1 |
| | CDC1551/ Oshkosh | Superoxide dismutase (Fe) P9WGE6] | 86 | |
| | ATCC25177/ H37Ra | Superoxide dismutase [A5U9H5] | 86 | |
| | ATCC25618/ H37Rv | Small ribosomal subunit protein uS7 [P9WH29] | 97.4 | |

| | | | | |
|---|---|---|---|---|
| 30S ribosomal protein S7 [A0A0J6Y7X7] | CDC1551/ Oshkosh | Small ribosomal subunit protein uS7 [P9WH28] | 97.4 | 30S ribosomal protein S7 OS=Mycolicibacterium obuense OX=1807 GN=rpsG PE=3 SV=1 |
| | ATCC25177/ H37Ra | Small ribosomal subunit protein uS7 [A5U069] | 97.4 | |
| Chaperonin GroEL [A0A0J6Y7E1] | ATCC25618/ H37Rv | Chaperonin GroEL 1 [P9WPE9] | 80.6 | Chaperonin GroEL OS=Mycolicibacterium obuense OX=1807 GN=groL2 PE=3 SV=1; Chaperonin GroEL OS=Mycolicibacterium obuense OX=1807 GN=groL1 PE=3 SV=1 |
| | CDC1551/ Oshkosh | Chaperonin GroEL 1 [P9WPE8] | 80.6 | |
| | ATCC25177/ H37Ra | Chaperonin GroEL 1 [A5U892] | 80.6 | |

**Supplemental Table 9: Sequence similarity between select *M. obuense* proteins and Mtb proteins.**

| MOS protein [Uniprot ID] | Top 3 hits after performing BLAST analysis on Mtb proteome | | |
|---|---|---|---|
| | Mtb strain | Protein name [Uniprot ID] | % Similarity |
| ATP synthase subunit beta [A0A0J6WFN0, A0A0M2K9M4] | ATCC 25618/ H37Rv | ATP synthase subunit beta [P9WPU5] | 92.5 |
| | CDC1551/ Oshkosh | ATP synthase subunit beta [P9WPU4] | 92.5 |
| | ATCC 25177/ H37Ra | ATP synthase subunit beta [A5U209] | 92.5 |
| Membrane protein [A0A0J6VFI3] | ATCC35801/ Erdman | Thioredoxin-like fold domain containing proteins [A0A0H3LHI5] | 55.3 |
| | ATCC25618/ H37Rv | Possible conserved membrane or secreted protein [O33272] | 55.3 |
| | ATCC25177/ H37Ra | Conserved membrane protein [A5U6X8] | 55.3 |
| UPF0182 protein [A0A0J6WAY1, A0A0M2JYV6] | T46 | UPF0182 protein [A0A7U8TRC8] | 79.3 |
| | CPHL_A | UPF0182 protein [A0A7U8TU30] | 79.3 |
| | ATCC25177/ H37Ra | UPF0182 protein [A5U7L0] | 79.3 |
| ATP synthase subunit alpha [A0A0J6WI07] | ATCC25618/ H37Rv | ATP synthase subunit alpha [P9WPU7] | 82.4 |
| | CDC1551/ Oshkosh | ATP synthase subunit alpha [P9WPU6] | 82.4 |
| | ATCC25177/ H37Ra | ATP synthase subunit alpha [A5U207] | 82.4 |
| Chaperonin GroEL [A0A0J6Y7E1] | ATCC25618/ H37Rv | Chaperonin GroEL 1 [P9WPE9] | 80.6 |
| | CDC1551/ Oshkosh | Chaperonin GroEL 1 [P9WPE8] | 80.6 |
| | ATCC25177/ H37Ra | Chaperonin GroEL 1 [A5U892] | 80.6 |
| Alanine and protein rich secreted protein [A0A0J6Z5J6] | ATCC25618/ H37Rv | Alanine and protein rich secreted protein (Apa) [P9WIR7] | 59.2 |
| | CDC1551/ Oshkosh | Alanine and protein rich secreted protein (Apa) [P9WIR6] | 59.2 |
| | ATCC35801/Erdman | Alanine and protein rich secreted protein (Apa) [A0A0H3LAS2] | 59.2 |

**Supplemental Table 10: Fc Array detection reagents.**

| Reagent Name | Source | Concentration at detection |
|---|---|---|
| Mouse anti-human IgG1 Hinge-PE | Southern Biotech, 9052-09 | 0.65 µg/mL |
| Mouse anti-human IgG2 Fc-PE | Southern Biotech, 9070-09 | |
| Mouse anti-human IgG3 Hinge-PE | Southern Biotech, 9210-09 | |
| Mouse anti-human IgG4 Fc-PE | Southern Biotech, 9200-09 | |
| Mouse anti-human Fc-PE | Southern Biotech, 9040-09 | |
| Mouse anti-human IgM-PE | Southern Biotech, 9020-09 | |
| Goat anti-human IgA-PE | Southern Biotech, 2050-09 | |
| Anti-monkey IgG | Life Diagnostics Inc., 10C2-9 | |
| Anti-monkey IgM | Life Diagnostics Inc., 2C11-1-5 | |
| Anti-monkey IgA | Nonhuman Primate Reagent Resource, AB_2819305 | |
| Goat anti-mouse IgG Fc PE | ThermoFisher Scientific, 31861 | 0.5 µg/mL |
| FcγR2A H131 (Biotinylated) | Duke Protein Production facility | 0.65 µg/mL |
| FcγR2B (Biotinylated) | Duke Protein Production facility | |
| FcγR3A V158 (Biotinylated) | Duke Protein Production facility | |
| FcγR3B (Biotinylated) | Duke Protein Production facility | |
| FcαR (Biotinylated) | Duke Protein Production facility | |
